# Pathogenic tau in the mouse locus coeruleus produces noradrenergic hyperactivity and neuropsychiatric phenotypes reminiscent of early Alzheimer’s disease

**DOI:** 10.64898/2026.01.30.702923

**Authors:** Anu Korukonda, Harris E. Blankenship, Korey Kam, Dilpreet Kour, Claudia Espinosa-Garcia, Wooyoung Eric Jang, Upasna Srivastava, Bernard Mulvey, Margaret M. Tish, Ronit Witztum, Christina C. Ramelow, Brittany S. Pate, L Cameron Liles, Linyue Lu, Jake Atallah, Eugene Guo, Keri Martinowich, Amanda L. Sharpe, Andrew W. Varga, Srikant Rangaraju, Michael J. Beckstead, David Weinshenker

## Abstract

Alzheimer’s disease (AD), though defined as a cognitive disorder, often presents neuropsychiatric symptoms such as anxiety, depression, agitation and sleep disruptions years before the onset of frank memory impairment. An early pathological feature is the accumulation of hyperphosphorylated “pretangle” tau (pTau) in the locus coeruleus (LC), the brain’s primary source of norepinephrine (NE). While clinical studies link LC pTau burden to behavioral abnormalities, causal mechanisms remain unclear. We developed a translationally-relevant mouse model that recapitulates the ‘LC-first’ phenomenon using cell type-specific viral expression of pathogenic P364S mutant human tau in LC neurons. Three months post-infusion, pTau accumulation induced anxiety-and compulsive-like behaviors and reduced sleep spindles without altering overall sleep architecture. Consistent with the behavioral phenotypes, electrophysiological recordings revealed significant increases in spontaneous and evoked firing of LC neurons, accompanied by robust astrocytic reactivity with no apparent cell death. Transcriptomic analysis identified upregulation of *Hcn2* and downregulation of *Clic6*, suggesting changes in neuronal excitability. To further define molecular mechanisms, we developed a cell type-specific proteomics approach, which showed synaptic and metabolic alterations associated with LC-specific tau pathology. Early anxiety-like behaviors observed at 3 months diminished at later timepoints (6-9 months) and were replaced by anxiolytic characteristics. These findings demonstrate that pTau triggers phenotypes reflective of LC-NE hyperactivity in the early stages of AD pathogenesis, laying the foundation for the development of LC-based disease-modifying therapies to address neuropsychiatric manifestations.

## Introduction

Alzheimer’s disease (AD) accounts for over 60% of dementia cases and is projected to affect 13.8 million people in the United States by 2060^1^. With a rapidly aging population, AD imposes a substantial socioeconomic toll on patients and caregivers^1^. Pathological hallmarks, amyloid-β(Aβ) plaques and tau neurofibrillary tangles (NFTs), begin accumulating years before cognitive symptoms^2^. This stage is often characterized by neuropsychiatric symptoms (NPS), collectively defined as mild behavioral impairment (MBI), a syndrome of persistent behavioral and personality change across motivational, affective, impulse-control, social and perceptual domains that precedes cognitive decline and confers greater dementia risk^3,4^.

As the primary source of norepinephrine (NE) in the central nervous system, the locus coeruleus (LC) modulates affect, arousal, attention, stress responses, vigilance, and the sleep-wake cycle^5–8^. Importantly, one of the earliest and defining pathological features coinciding with NPS onset is the emergence of hyperphosphorylated pretangle tau (pTau) in the LC, termed Braak stage 0^9–11^. Prior to forebrain Aβ plaque pathology, pTau preferentially accumulates in somatodendritic compartments and proximal axons of LC neurons that densely innervate limbic and neocortical areas, structures that subsequently exhibit aberrant tau aggregation in the form of NFTs^9–12^. As disease progresses through Braak stages, LC tau pathology intensifies and is associated with ∼8.4% LC volume reduction per stage, with significant neuronal loss by Braak stage III and catastrophic degeneration by late stages of AD^13,14^. Among other monoaminergic nuclei, only LC cell density shows a significant correlation with cognitive function, highlighting the clinical relevance of early LC vulnerability^15^.

Early pTau accumulation in the isodendritic core (including LC) is associated with increased odds of agitation, depression, anxiety, and sleep disturbances during initial stages of the disease^16^. Longitudinal studies report that individuals with early-onset AD exhibit higher apathy and anxiety with worse LC structural integrity compared to late-onset AD^17,18^. Disruptions in LC integrity also correlate with daytime dysfunction, nocturnal awakenings, impaired episodic memory, and increased likelihood of developing mild cognitive impairment (MCI) ^19^. Elevated LC signal in cortical tau-positive subjects predicts NPS severity, suggesting compensatory escalations in NE transmission may promote impulsivity and aggression^20^. These data align with cerebrospinal fluid (CSF) and postmortem studies, where NE and its metabolites, 3-methoxy-4-hydroxyphenylglycol (MHPG) and its neurotoxic byproduct 3,4-dihydroxyphenylglycolaldehyde (DOPEGAL), are consistently higher in MCI/early AD, and linked to agitation and poor impulse control^17,21–24^. In parallel, LC neuron loss is accompanied by increased gene expression of the rate-limiting NE biosynthetic enzyme, tyrosine hydroxylase (TH), in surviving cells^25,26^. Microarray analyses of postmortem LC tissue from MCI cases also implicate disruptions in oxidative stress, mitochondrial function, cytoskeletal proteases, and axonal degeneration pathways^27^. Because tau pathology typically inhibits neuronal activity, a prevailing hypothesis is that pTau-bearing LC neurons initially engage homeostatic plasticity mechanisms and become hyperactive, thereby dysregulating NE signaling in limbic-and cortical-projecting circuits^25–29^. Limitations in detecting LC pTau in living patients and reliance on late-stage postmortem tissue constrain causal relationships between LC dysregulation and NPS.

Animal studies support a mechanistic role for LC-NE dysfunction, particularly in late-stage LC degeneration. Partial LC lesions or genetic/chemical NE depletion in transgenic AD mouse models exacerbates AD-like neuropathology and cognitive impairment, whereas enhancing LC-NE transmission ameliorates such deficits^24,30–37^. Yet efforts linking LC tau pathology to noradrenergic dysregulation and behavioral consequences have been limited by (i) ubiquitous transgene promoters driving pTau accumulation outside the LC and failing to recapitulate the “LC first” pattern, (ii) presence of Aβ pathology from overexpression of mutant amyloid genes, (iii) failure to measure LC firing, (iv) insufficient behavioral paradigms capturing NPS phenotypes, and/or (v) incomplete characterization of LC transcriptomic and proteomic adaptations to tau inclusions^38–44^. We thus sought to develop a biologically-relevant model that restricts pTau to the LC to test causal mechanisms during the earliest AD stages, when disease-modifying interventions may be most effective.

To address this gap, we combined TH-Cre mice with a Cre-dependent viral vector to overexpress a mutant form of human tau (P364S) specifically in the LC. This variant causes frontotemporal dementia (FTD) in humans and confers susceptibility to hyperphosphorylation and aggregation while reducing microtubule assembly function^45,46^. We chose P364S over canonical P301S/L mutations because it lacks seeding capacity *in vitro*, reducing likelihood of propagation to LC-connected regions and strengthening LC-intrinsic interpretation of downstream effects^47^. Neuropathology and behavior were assessed at 3, 6, and 9 months post-infusion, timepoints based on previously reported LC pTau progression^33^. In addition, we performed slice electrophysiology as well as cell-type-specific transcriptomics and proteomics at the earliest timepoint to identify putative LC-intrinsic cellular and molecular alterations induced by tau inclusions and downstream pathways contributing to dysregulated LC activity and NPS phenotypes.

## Materials and Methods

### Animal studies

Adult male and female mice (2-3 months) were used. TH-Cre mice (B6.Cg-7630403G23Rik^Tg(Th-cre)1Tmd^/J, The Jackson Laboratory, #008601) were utilized for behavior, immunofluorescence, and slice electrophysiology. For transcriptomics, TH-Cre mice were crossed with Slc6a2-eGFP/Rpl10a mice (B6;FVB-Tg(Slc6a2-eGFP/Rpl10a)JD1538Htz/J, Jackson Laboratory, #031151), which integrates the fusion ribosomal protein, EGFP/Rpl10a, into a bacterial artificial chromosome under the Slc6a2 (NE transporter) promoter^48^. For proteomics, TH-Cre mice were crossed with Rosa26^TurboID/WT^ mice (C57BL/6-Gt(ROSA)26Sor^tm1(birA)Srgj^/J, Jackson Laboratory, #037890) ^49^. Mice were group-housed with age-and sex-matched conspecifics until 1 week prior to behavioral testing, then singly-housed until tissue collection. Mice were maintained on a 12 h light/dark cycle with *ad libitum* access to food and water. All experiments were conducted in the vivarium at Emory University, the Oklahoma Medical Research Foundation, and the Icahn School of Medicine in accordance with the NIH Guideline for the Care and Use of Laboratory Animals and approved by the respective Institutional Animal Care and Use Committees.

### Viral Vectors

An AAV5-EF1α-DIO-P364S human tau construct was developed by the Emory Custom Cloning and Viral Vector Cores. The commercially available AAV5-EF1α-DIO-EYFP virus served as control (Addgene, plasmid #27056).

### Stereotaxic surgeries

Mice were anesthetized with 3% isoflurane and administered ketoprofen (5 mg/kg, s.c.). AAV5-EF1α-DIO-P364S human tau or AAV5-EF1α-DIO-EYFP was infused (500 nL, 150 μL/min) into the mouse LC bilaterally at the following coordinates (AP:-5.4 mm, ML: ±1.2 mm, DV:-4.0 mm relative to bregma). For time course assessments of tau pathology, unilateral LC infusions of AAV5-EF1α-DIO-P364S was performed. Following a 5 min post-infusion period, the needle was withdrawn, the scalp was closed with cyanoacrylate (3M Vetbond), and mice recovered on a heating pad. For sleep studies, mice underwent a second surgery for implantation of EEG/EMG electrodes 2-3 weeks prior to sleep studies as previously described ^50^.

### Behavioral paradigms and sleep studies

NE-sensitive behavioral assays were conducted from least to most stressful during the light cycle^51–53^. Testing comprised of nestlet shredding (NS) and marble burying (MB) to evaluate compulsive-like behaviors, novelty-suppressed feeding (NSF) to assess for anxiety-like behavior, and fear conditioning (FC) to examine associative memory. See Supplementary Methods for details.

### Tissue collection

Mice were euthanized by overdose of sodium pentobarbital (Fatal Plus, 150 mg/kg, i.p.; Med-Vet International, Mettawa, IL) and transcardially perfused with ice-cold 4% PFA or PBS for immunofluorescence (IF) or proteomics, respectively. For IF, brains were post-fixed overnight in 4% PFA at 4°C and cryoprotected in 30% sucrose prior to sectioning. For proteomics, the hindbrain was dissected (cerebellum discarded), flash-frozen on dry ice, and stored in-80°C until processing.

For electrophysiology and translating ribosome affinity purification experiments, mice were anesthetized with isoflurane and euthanized by rapid decapitation. For electrophysiology, brains were submerged in an ice-cold cutting solution containing (in mM) 110 choline chloride, 2.5 KCl, 1.25Na2PO4, 0.5CaCl2, 10MgSO4, 25 glucose, 11.6 Na-ascorbate, 3.1 Na-pyruvate, 26 NaHC03, 12 N-acetyl-L-cysteine, and 2 kynurenic acid. For TRAP, the pons was rapidly dissected on ice, flash-frozen in dry ice, weighed, and stored at-80°C until processing.

### Immunofluorescence (IF) and microscopy

Forty μm thick coronal sections were collected, and IF was performed as previously described^32^. Antibody details are provided in Table 1 in Supplementary Methods. All images were acquired on the Keyence BZ-X700 microscope system as z-stacks (10 z-stacks, pitch: 1 µm) at 2x or 20x magnification. Full focus feature was used for image compression. One representative atlas-matched section was selected per animal for the three regions: LC, dorsal HC, and PFC. For TH^+^ cell counts, all sections containing the LC in each animal were imaged. Image processing and quantification were conducted on FIJI/ImageJ software. The pipeline included background subtraction, intensity thresholding for each marker within region of interest and integrated density measurements (for glial cell cells, size = 5-Infinity µm). The Cell Counter plugin was used for counting TH^+^ cells manually.

### Electrophysiology

For more details on slice preparation and recordings, see Supplementary Methods.

### Translating Ribosome Affinity Purification (TRAP) and RNA sequencing

Samples from two age-, sex-, and treatment-matched TH-Cre+, Slc6a2-eGFP/Rpl10a+ mice were pooled to form a biological replicate. TRAP was performed as described to enrich for LC ribosomes^48^. RNA was extracted, quality-controlled, and libraries prepared using SMART-Seq v4 and Nextera XT kits. Sequencing was performed on an Illumina NovaSeq X Plus (150 bp PE). Reads were aligned to mm10 and human tau using STAR; counts were generated with featureCounts. Sequencing data are available on NCBI GEO (GSE314950). Full bioinformatic pipeline on normalization parameters (model: *∼0+transgene+sex+TRAPefficiency*), differentially expressed genes (DEGs) analysis and Gene Set Enrichment Analysis (GSEA) are detailed in the Supplementary Methods.

### Cell type-specific *in vivo* biotinylation of proteins (CIBOP)

TH-Cre;*Rosa26^TurboID/WT^* mice and their littermates (WT or TH-Cre) were given water supplemented with biotin (37.5 mg/L) for 2 weeks prior to euthanasia. Homogenization of frozen tissue and quantification of total protein concentration were conducted as previously described^49^. Biotinylation and enrichment were validated by immunoblotting. Biotinylated proteins were affinity-purified using streptavidin magnetic beads following established protocols^49^. Samples were analyzed by DIA LC–MS/MS using a Vanquish Neo UHPLC coupled to an Orbitrap Exploris 480. Data are available via PRIDE (PXD071656). Full MS methods and analysis pipelines including data processing, quantification, normalization parameters, differentially expressed proteins (DEPs), and Gene Ontology (GO) and network enrichment analysis are provided in Supplementary Methods.

## Statistical analyses

All data are reported as mean ± SEM with statistical significance set at p < 0.05. Behavioral, IF and electrophysiology data were compared between EYFP and Tau groups using an unpaired t-test (or Welch t-test when variance assumptions were violated) or Mann-Whitney U test, as determined in GraphPad Prism 10. For behavioral assessments such as NS, NSF, and FC, a two-way ANOVA was used with factors such as treatment, location/time where appropriate. Outliers were identified and removed with Grubb’s test. For statistical details related to omics analyses, see corresponding sections.

### Viral-mediated expression of mutant P364S human tau in the LC triggers pTau at 3 months

Since pTau emerges in the human LC decades before cognitive deficits and coincides with MBI^9,10,16^, we developed a viral approach to selectively express pTau in noradrenergic neurons. 2-3 month old TH-Cre mice received intra-LC infusions of AAV5-DIO-P364S human tau (Tau) or control AAV5-DIO-EYFP (EYFP) and were assessed 3 months later **(Fig. 1A)**. Robust bilateral expression of human tau or EYFP were confirmed within TH^+^ LC neurons **(Fig. 1B, C)**. Quantification of pTau Ser202/Thr205 using AT8 immunoreactivity (IR) showed a significant treatment effect with Tau virus inducing pTau in somatodendritic compartments of LC neurons (t_(5.1)_ = 2.692, p < 0.05, Welch’s correction) **(Fig. 1D, E)**.

**Figure 1.**
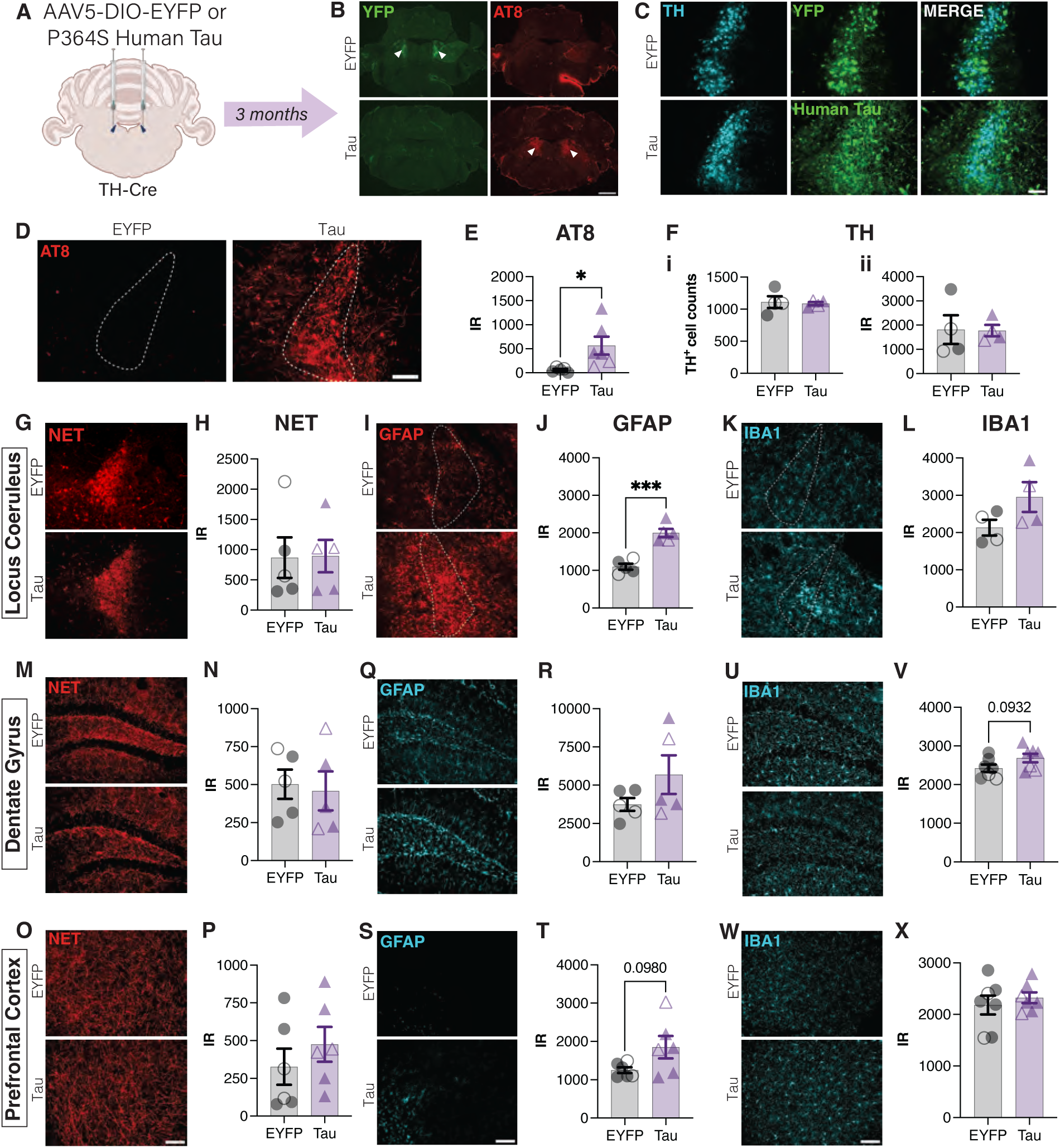
Cell-type specific expression of P364S mutant human tau at 3 months triggers hyperphosphorylated tau accumulation and astrogliosis in the mouse LC. **(A)** TH-Cre mice received bilateral LC infusions of AAV5-DIO-EYFP (EYFP) or P364S mutant human tau (Tau). Three months later, mice were evaluated for LC integrity and pathology. **(B)** Verification of site-specificity of YFP (green) and AT8 (red) in the LC (depicted by white arrows, 2x, scale bar = 1000 μm) in the EYFP and Tau mice, respectively. **(C)** Colocalization of YFP (green) and human tau (green) expression in the LC, delineated by TH (cyan). Representative images and quantification of immunoreactivity (IR) of **(D, E)** AT8 (red), **(F)** TH (cyan), **(G, H)** NET (red), **(I, J)** GFAP (red), and **(K, L)** IBA1 (cyan) (LC outlined in white; 20x, scale bar = 100 μm). Representative IF images and quantification of IR in the DG and PFC for the following markers **(M-P)** NET (red), **(Q-T)** GFAP (cyan), and **(U-X)** IBA1 (cyan) (20x; scale bar: 100 μm). Data are presented as mean ± SEM (n = 5-6 per group; male: closed, female: open symbols). Unpaired t-test with Welch’s correction where applicable. *p < 0.05, ***p < 0.001.

To examine whether tau pathology impacted LC integrity, we quantified TH^+^ cells and TH and NE transporter (NET) IR across the rostrocaudal axis, all of which were unchanged **(Fig. 1F-H)**. In terms of local neuroinflammatory responses, pTau significantly increased GFAP IR (t_(8)_ = 6.730, p < 0.0005), but not IBA1 IR **(Fig. 1I-L)**. Together, these data demonstrate that P364S tau provokes robust hyperphosphorylation and an astrocytic response without overt LC neuronal loss.

### LC tau pathology does not affect NE terminal density at 3 months

To test the “dying-back” phenomenon, in which LC projections degenerate before cell body loss^24,34^, we examined LC target regions with dense noradrenergic innervation, such as the dentate gyrus (DG) of hippocampus and prefrontal cortex (PFC). NET IR revealed no treatment differences in noradrenergic fiber density in either region at 3 months **(Fig. 1M-P)**. Astrocytic GFAP IR was unchanged in the DG but showed an upward trend in the PFC **(Fig. 1Q-T)**. Conversely, microglial IBA1 IR was comparable across groups in the PFC, but trended higher in the DG of Tau-expressing mice **(Fig. 1U-X)**.

### pTau drives anxiogenic behaviors consistent with LC hyperactivity at 3 months

Early AD/MBI is associated with elevated CSF NE and NPS consistent with LC-NE hyperactivity^17,24^. We previously showed that chemogenetic, optogenetic, or pharmacological activation of LC-NE signaling promotes compulsivity, anxiety-like behavior, and wakefulness^51,54,55^, and that pathogenic LC inclusions or perturbations to NE transmission in rodent AD and Parkinson’s disease (PD) models elicit anxiogenic behaviors^32,39,56^. We thus evaluated consequences of pathogenic tau expression on NE-sensitive behaviors at 3 months post-infusions.

For NS, a measure of stress-induced compulsive-like behavior, a two-way ANOVA revealed a main effect of treatment (F_(1,20)_ = 4.457, p < 0.05), with Tau-expressing mice displaying increased shredding at 2 and 4 h following cage change **(Fig. 2A)**. Anxiety-like and neophobic behavior were assessed using the NSF task, with a home-cage control to account for satiety^52^. There were main effects of treatment (F_(1,18)_ = 6.16, p < 0.05) and environment (F_(1, 36)_ = 45.61, p < 0.0001), and a treatment × environment interaction (F_(1, 36)_ = 7.324, p < 0.05). Post-hoc Sidak’s comparison revealed that Tau-expressing mice exhibited longer latencies to consume food pellets in the novel arena compared to EYFP (p < 0.005), while home-cage latencies were unchanged, indicating novelty-induced anxiety rather than altered appetite **(Fig. 2B)**.

**Figure 2.**
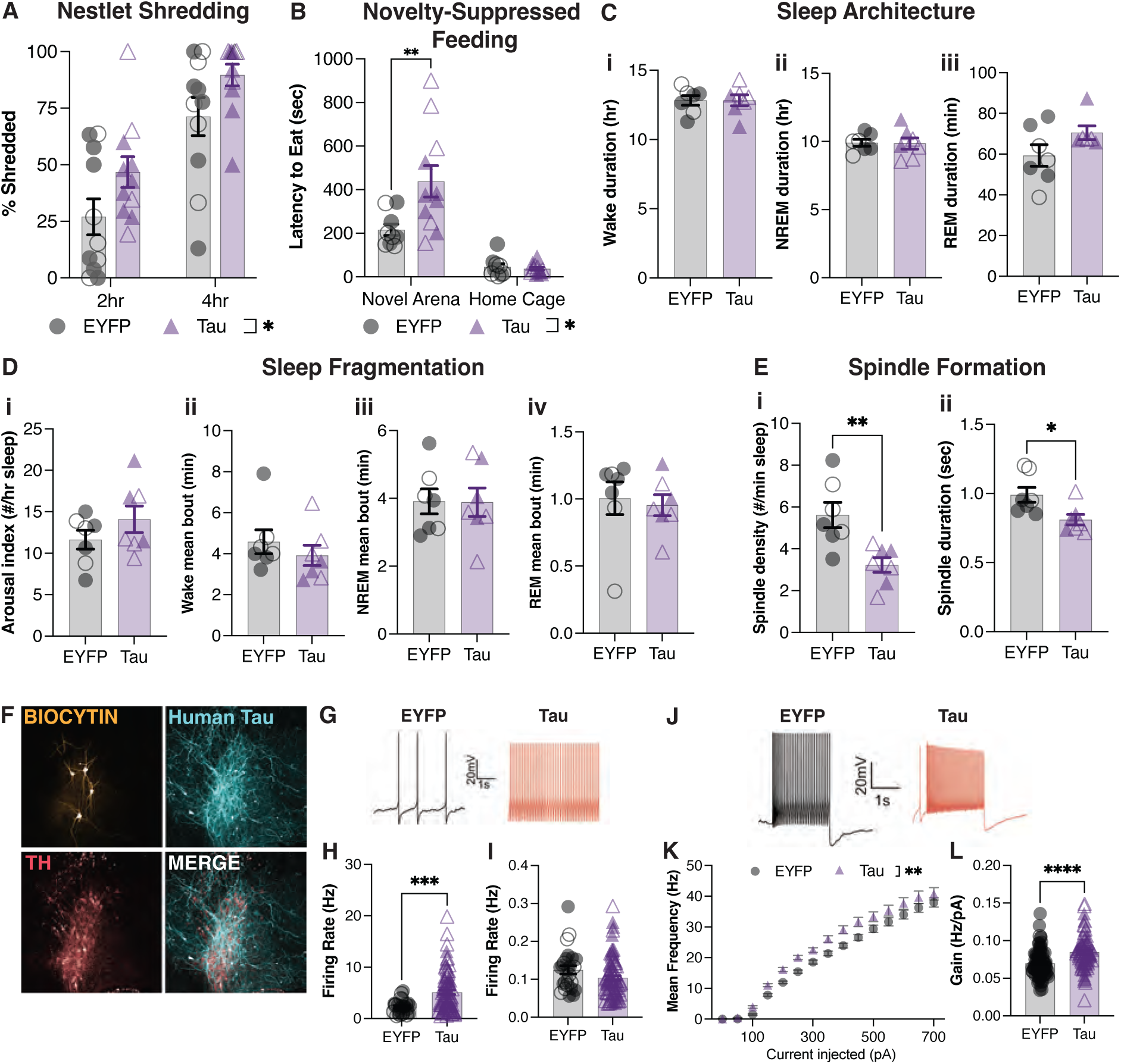
Hyperphosphorylated tau drives anxiety-like behaviors, reduces sleep spindles, and provokes LC hyperactivity at 3 months. TH-Cre mice received bilateral LC infusions of EYFP or Tau. Three months later, mice were evaluated for behavior, EEG sleep parameters, and LC neuronal activity. **(A)** Bar graph showing the percent of nestlet shredded at 2 and 4 h following cage change (n = 10-11 per group, two-way ANOVA). **(B)** Bar graph displaying latency to eat in the novel arena and home cages (n = 10-11 per group, 2-way ANOVA with Sidak’s post hoc test). **(C)** Mean durations for (i) wake, (ii) NREM, and (iii) REM to assess for sleep architecture. (**D)** Bar graphs of arousal index (i) and mean bouts for (ii) wake, (iii) NREM, and (iv) REM to evaluate sleep fragmentation. **(E)** Sleep density (i) and spindle duration (ii) recorded to characterize sleep microstructures (C-E: n = 7 per group, unpaired t-test). **(F)** Representative images of biocytin-filled neurons (yellow), human tau (cyan), and TH (red) in the LC of mice injected with Tau. **(G)** Representative traces of spontaneous firing in EYFP (black) and Tau (red) LC-NE neurons (x-axis: 1 sec, y-axis: 200 mV). **(H, I)** Bar graphs for spontaneous firing rate and the coefficient of variation (CV) of the interspike interval of LC-NE neurons (Mann-Whitney U test). **(J)** Representative traces displaying 2 sec long depolarization steps to 100pA in EYFP and Tau LC-NE neurons (x-axis: 1 sec, y-axis: 200 mV). **(K)** Mean evoked spike frequency as a function of injected current (1-750 pA; two-way ANOVA test). **(L)** Gain (25-125 pA) quantified as slope of the frequency-current (F-I) curve (Mann-Whitney U test). (G-L: n = 25-84 recordings per group; male: closed, female: open symbols). Data are presented as mean ± SEM. *p < 0.05, ** p < 0.01, ***p < 0.001, ****p < 0.0001.

### LC tau pathology affects sleep spindles without disrupting sleep architecture at 3 months

Because the LC-NE system influences wakefulness^6,57^, we determined if aberrant tau affected the sleep-wake cycle. Sleep macrostructure was intact, with no differences in wake, NREM, or REM durations **(Fig. 2C)**. Sleep fragmentation, measured by arousal index and bout length, was also unperturbed **(Fig. 2D)**. However, Tau-expressing mice displayed significant reductions in NREM spindle density (t_(12)_ = 3.43, p = 0.005) and duration (t_(12)_ = 2.71, p < 0.05), consistent with LC hyperactivity^58,59^ **(Fig. 2E)**.

### Tau expression increases spontaneous and evoked LC activity at 3 months

To directly test whether pTau alters LC firing, we performed *ex vivo* slice electrophysiology. Putative noradrenergic cells were identified electrophysiologically and confirmed post-hoc by biocytin filling and human tau expression **(Fig. 2F)**. Spontaneous firing rates were elevated in Tau-expressing LC neurons (U = 558, p < 0.0005) without disrupting spike rhythmicity **(Fig. 2G-I)**. For evoked firing, a two-way ANOVA showed main effects of treatment (F_(1,166)_ = 8.21, p < 0.005) and current injection (F_(14,2310)_ = 580.9, p < 0.0001). Quantification of the frequency-current curve showed a steeper steady-state slope in Tau neurons compared with controls (U = 1790.5, p < 0.0001), indicating increased neuronal excitability **(Fig. 2J-L)**.

### Tau-induced changes to the LC translatome at 3 months

To define LC-intrinsic molecular compensatory adaptations to tau pathology, we conducted TRAP to isolate EGFP-tagged ribosomes selectively from noradrenergic neurons using NET gene (*Slc6a2*) promoter-driven expression **(Fig. 3A)**. Sequencing confirmed the presence of human P364S tau transcripts in Tau-treated LCs and absence in EYFP controls **(Suppl. Table 1)**. Validation of TRAP enrichment using paired tissue homogenates and TRAP immunoprecipitated (IP) fractions demonstrated robust enrichment of canonical LC markers (*Slc6a2*, *Th*, *Dbh*) in both groups, with higher enrichment efficiency in Tau samples **(Suppl. Fig. 1)** ^48,60^.

**Figure 3.**
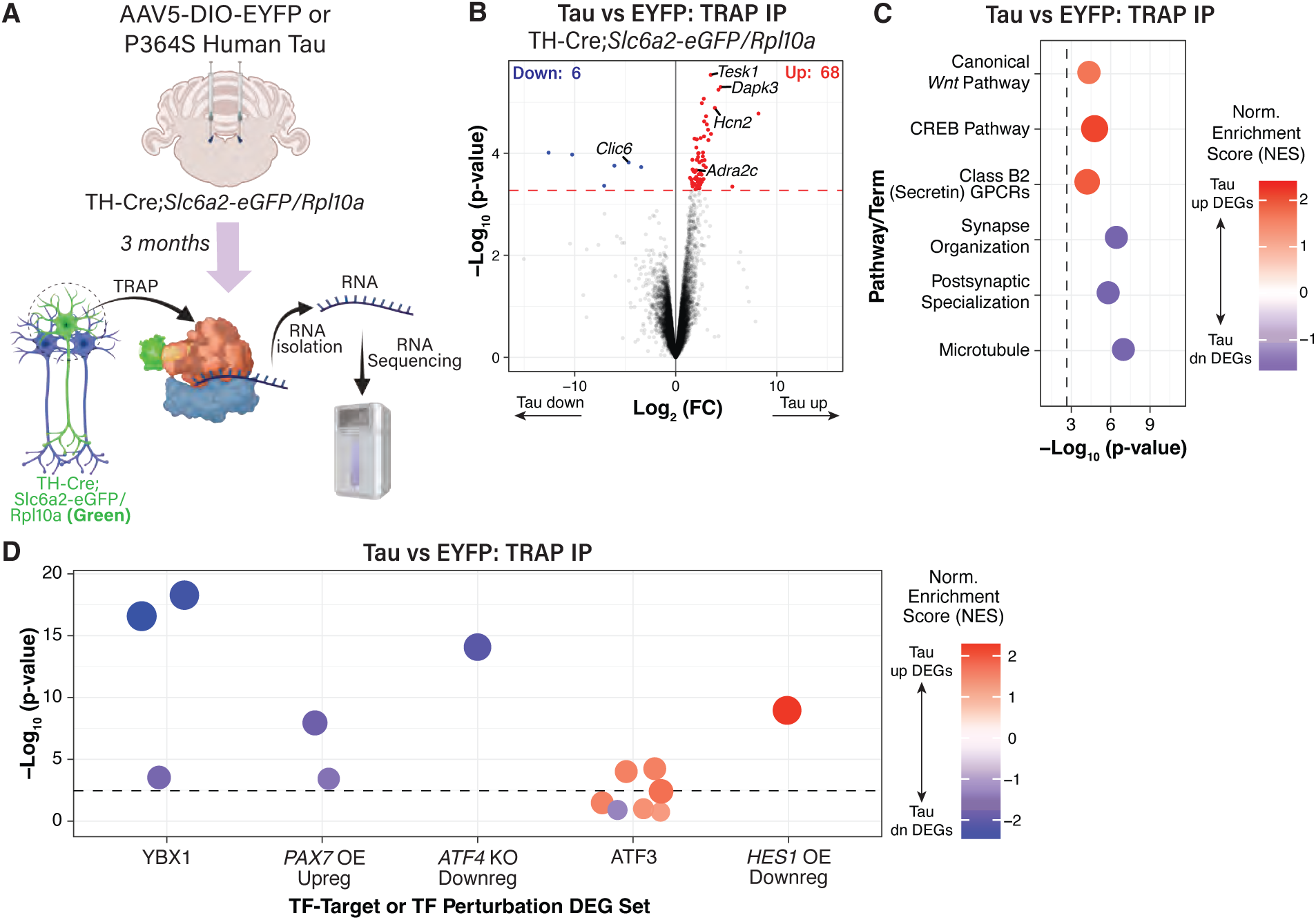
Tau induces alterations to the LC translatome at 3 months. **(A)** TH-Cre;*Slc6a2-eGFP/Rpl10* mice received bilateral LC infusions of EYFP or Tau. Three months later, tissue was harvested and TRAP was performed to isolate EGFP-Rpl10a-tagged ribosome-bound mRNA from LC neurons. **(B)** Volcano plot depicting DEGs comparing Tau and EYFP TRAP IP fractions, adjusting for sex and TRAP enrichment efficiency. The red line denotes the FDR < 0.1 threshold for DEG significance. **(C)** Pathway and ontology enrichment of Tau DEGs. Selected results from GSEA using MSigDB gene sets are illustrated. Positive normalized enrichment score (NES) signifies enrichment among Tau-upregulated genes, and negative NES signifies enrichment in Tau-downregulated genes. The dashed line indicates FDR < 0.05. **(D)** Enrichment of Tau DEGs transcription factor (TF) regulatory targets. Selected results from GSEA using TF-target gene sets from Enrichr (OE: overexpression; KO: knockout). Positive NES signifies enrichment in Tau-upregulated genes, and negative NES signifies enrichment in Tau-downregulated genes. Dashed line indicates FDR < 0.05 (n = 3-4 per group).

Differential expression analysis of 14,224 genes, controlling for condition-dependent TRAP efficiency (see Supplementary Methods), identified 74 DEGs at FDR < 0.1, the majority of which were upregulated in Tau-expressing mice **(Fig. 3B, Suppl. Table 1)**. Notably, reciprocal regulation of excitability-related genes was observed, including downregulation of the inhibitory chloride channel *Clic6* and upregulation of the excitatory pacemaker channel *Hcn2* **(Fig. 3B)**. The adrenergic receptor 𝛼_2C_ (*Adra2c*) was also upregulated, suggesting compensatory responses to LC-NE hyperactivity **(Fig. 3B)**.

GSEA using MsigDB pathways revealed that Tau-upregulated genes were enriched in CREB, Wnt, and Secretin GPCR signaling, whereas downregulated genes were enriched in synaptic compartments **(Fig. 3C, Suppl. Table 1)**, indicating coordinated synaptic plasticity alterations^61,62^. GSEA of transcription factor target gene sets identified enrichment of Atf3 overexpression-induced genes among Tau-upregulated DEGs and repression of Ybx1-and Atf4-dependent genes **(Fig. 3D; Suppl. Table 1)**. Tau-downregulated genes were also enriched for Pax7-dependent targets, consistent with altered expression of LC identity programs^63^, and validating our TRAP efficiency normalization, as unadjusted analyses would spuriously reduce Pax7 targets in lower-enriched EYFP samples. Genes repressed by Hes1 were also overrepresented among upregulated DEGs, suggesting reduced Hes1 activity. Together, these findings indicate a shift in proteostasis regulation from Atf4 toward Atf3, de-repression of Hes1 targets, and potential loss of pontine identity genes downstream of Pax7 **(Fig. 3D; Suppl. Table 1)**.

### Noradrenergic-specific proteomic profiling reveals Tau-induced alterations in the LC at 3 months

Because transcriptomic and proteomic changes often diverge due to post-transcriptional and post-translational regulation^64^, we examined pTau-induced proteomic alterations in LC neurons. Given the limited spatial resolution of bulk proteomics for small nuclei such as the LC (∼3,000 neurons), we employed CIBOP, a proximity-based labeling approach^49^. TurboID expression was restricted to catecholaminergic neurons by crossing *Rosa26^TurboID/WT^* mice with TH-Cre animals. Brainstem dissections excluding midbrain dopaminergic regions showed robust biotin labeling in TH^+^ neurons after two weeks of biotin supplementation, confirming cell-type specificity and feasibility **(Suppl. Fig. 2A, B)**.

Three months following intra-LC infusion of EYFP or P364S human tau into TH-Cre; *Rosa26^TurboID/WT^*mice, brainstems were processed using an optimized CIBOP workflow^49,65^ **(Fig 4A)**. IF and WB confirmed robust biotinylation in TH^+^ soma and dendrites **(Fig. 4B, C)**, and expression of pTau231 in Tau-treated samples **(Fig. 4C)**. Streptavidin-based enrichment of biotinylated proteins (pulldown) was validated by WB **(Fig. 4D)**. MS identified 6,666 proteins in input samples and 4,579 proteins in the streptavidin-enriched fraction. Pulldowns were enriched for canonical LC markers (TH, DBH, NET), confirming successful cell-type specificity **(Fig. 4E)**.

**Figure 4.**
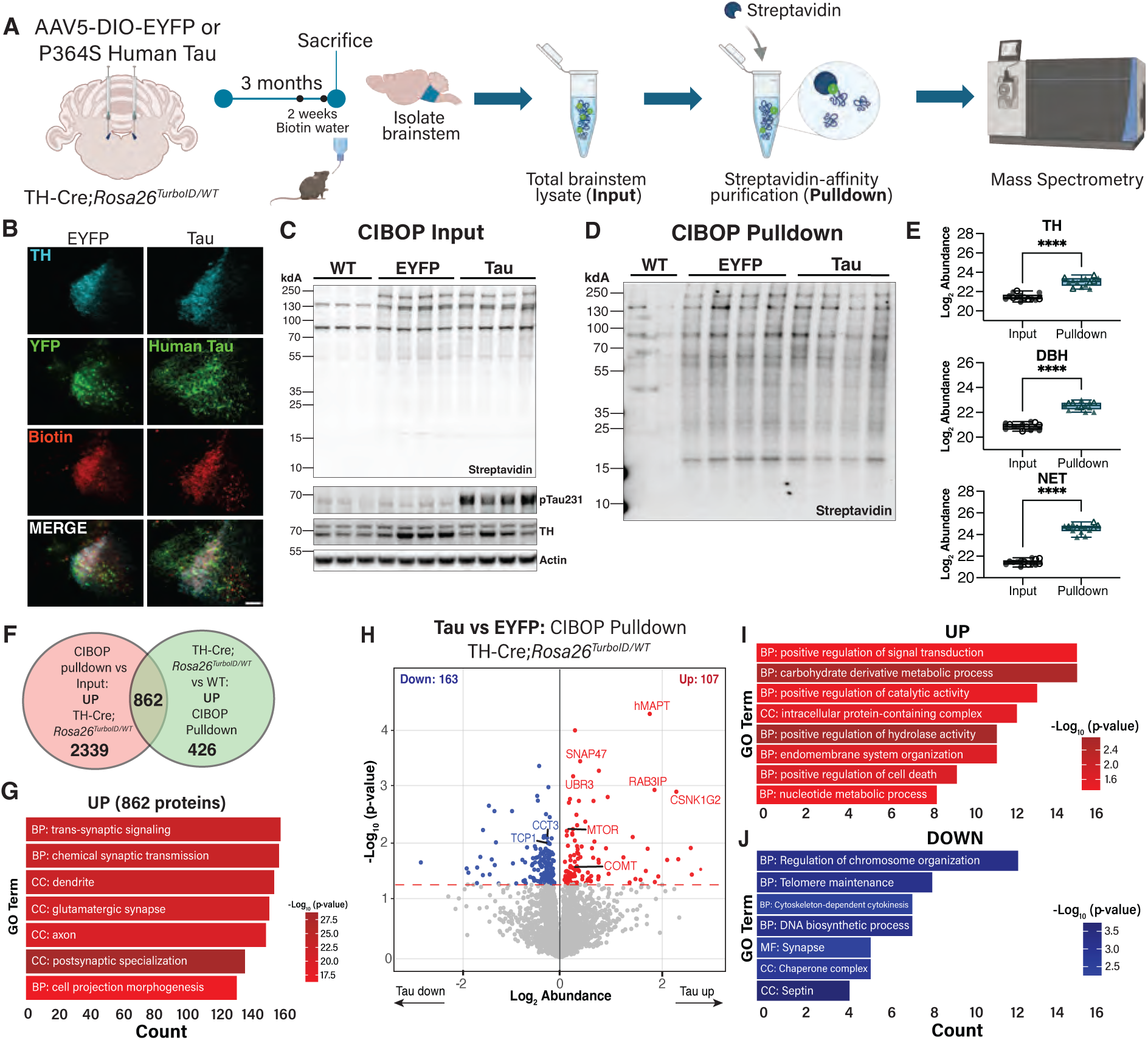
CIBOP for LC-NE proteome reveals modest changes in response to pathogenic tau at 3 months. **(A)** Schematic of the study design for LC-specific proteomic biotinylation using CIBOP. **(B)** Representative IF images showing colocalization of YFP or human tau (green) with biotin labeling (red) in TH^+^ cells (cyan) (20x, scale bar = 100μm). **(C)** Representative western blots from brainstem lysates (WT, EYFP, and Tau) probed for biotin (streptavidin), TH, pTau-T231, and actin. **(D)** Western blot of enriched biotinylated proteins following streptavidin-affinity pulldown. **(E)** Boxplots depicting enrichment of canonical LC markers (TH, DBH, and NET) in the CIBOP pulldown compared to input (****p < 0.0001). **(F)** Venn diagram comparing proteins enriched in the CIBOP pulldown vs input fraction in TH-Cre;*Rosa26^TurboID/WT^* and proteins enriched in the CIBOP pulldown fraction of TH-Cre;*Rosa26^TurboID/WT^* vs WT at nominal p < 0.05. **(G)** GO pathway enrichment for the 862 proteins shared between the two comparisons shown in (F). **(H)** Volcano plot showing DEPs comparing Tau and EYFP in the CIBOP pulldown fraction. The red line denotes the p < 0.05 for DEP significance. **(I, J)** GO pathway enrichment of Tau DEPs (n = 7 per group).

To define a high-confidence LC proteome, we intersected proteins enriched in (i) CIBOP pulldown vs input in TH-Cre;*Rosa26^TurboID/WT^*mice and (ii) TH-Cre;*Rosa26^TurboID/WT^* vs WT pulldowns, yielding 862 LC-enriched proteins **(Fig. 4F)**. GO analysis revealed enrichment for postsynaptic specialization, glutamatergic synapses, and cell projection morphogenesis, along with canonical LC-NE markers (DBH, TH, NET, DDC, VMAT) **(Fig. 4G; Suppl. Table 2)**.

Next, we examined pathogenic tau-induced proteomic alterations. As validation, human MAPT was robustly enriched in both input (7-fold) and pulldown (9-fold) **(Suppl. Table 3, Fig. 4H)**. Within the streptavidin-enriched proteome, 270 DEPs (UP: 107, DOWN: 163; p < 0.05) were identified, including increased vesicle and synaptic trafficking regulators (SNAP47 and RAB3IP), catechol-O-methyltransferase (COMT), which metabolizes catecholamines including NE^66^, ubiquitin ligase (UBR3), and mTOR, which is upregulated in AD^67,68^, alongside reduced TCP1/CCT chaperonin complex components involved in proteostasis **(Fig. 4H)**. GO analysis of Tau-UP proteins was associated with metabolic pathways, intracellular signaling, and enzymatic activity **(Fig. 4I)**. Tau-DOWN proteins were enriched for genome maintenance, cytoskeletal, and synaptic scaffolding pathways, consistent with a cellular stress profile **(Fig. 4J)**. LC Tau-regulated proteins mapped selectively to specific human AD brain co-expression modules, with Tau-UP proteins enriched in M8 (intracellular trafficking/protein transport) and Tau-DOWN proteins enriched in M36 (synaptic/neurotransmitter modulation), indicating direction-dependent network engagement by Tau **(Suppl. Fig. 2C)**.

### Tau pathology, but not anxiogenic phenotypes, persists at 6 and 9 months

To model later disease stages, mice were aged to 6 or 9 months following viral infusions **(Fig. 5A, J)**. Expression of EYFP and human tau persisted at both timepoints **(Fig. 5B, K)**. Prolonged Tau exposure did not alter TH^+^ cell number; however, TH IR was significantly reduced at 6 months (t_(12)_ = 2.62, p < 0.05), with a similar, nonsignificant trend at 9 months, likely reflecting variability in viral expression **(Fig. 5C, L)**. AT8 IR was robust at both 6 months (t_(11)_ = 11.34, p < 0.0001) and 9 months (t_(6.0)_ = 8.45, p = 0.0001; Welch’s correction) **(Fig. 5D, E, M, N)**. NET IR and neuroinflammatory markers in the LC were unchanged at either timepoint **(Suppl. Fig. 3A-L)**, and NET signal in the DG or PFC also remained stable **(Fig. 5F-I, O-R)**.

**Figure 5.**
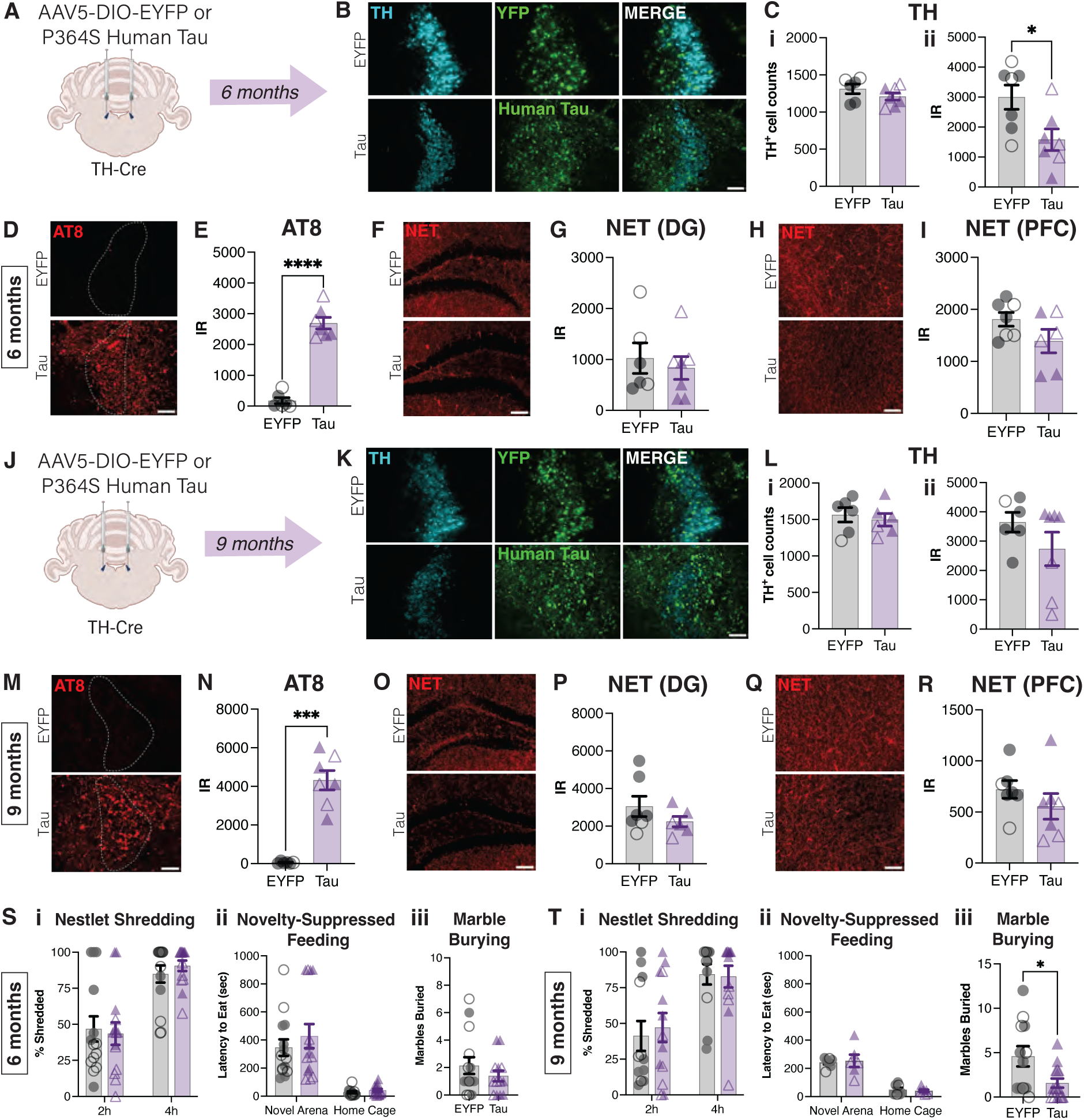
Prolonged pathogenic tau burden reverses anxiety-like behavior. Mice received stereotaxic infusion of EYFP or Tau into the LC and were evaluated for LC integrity, tau pathology, and behavior at **(A)** 6 months and **(J)** 9 months. **(B, K)** Colocalization of YFP (green) and human tau (green) expression in the LC, delineated by TH (cyan). Representative IF images and quantification of IR of **(C, L)** TH **(D, E, M, N)** AT8 (red; LC outlined in white), **(F-I, O-R)** NET (red) in the DG and PFC (n = 6-7 per group; 20x scale bar = 100 μm; unpaired t-test). **(S, T)** Behavioral analyses at 6 and 9 months showing (i) percent of nestlet shredded at 2 and 4 h following cage change, (ii) latency to eat in novel arena and home cage, and (iii) number of marbles buried (n = 12-14 per group; i and ii: two-way ANOVA, iii: unpaired t-test). Data are presented as mean ± SEM (male: closed, female: open symbols). *p < 0.05, ***p < 0.0001.

To assess potential escalation of tau pathology, we performed unilateral LC infusions of P364S human tau and immunostained with AT8 and PHF1, which detects more advanced tau species (pS396/pS404). Within-subject comparisons revealed minimal IR in control hemispheres and significant treatment effects for both AT8 (F_(1,9)_ = 54.28, p < 0.0001) and PHF1 (F_(1,10)_ = 20.78, p = 0.001), with no age-dependent differences **(Suppl. Fig. 2M-P)**. pTau propagation to LC projection regions was negligible, with little to no AT8 IR in the DG or PFC at any timepoint **(Suppl. Fig. 3Q)**.

In contrast to 3-month timepoint, anxiogenic and compulsive-like phenotypes were absent or reversed at later timepoints. At 9 months, Tau-expressing mice buried fewer marbles than controls (t_(15.3)_ = 2.42, p < 0.05; Welch’s correction), consistent with an anxiolytic shift, whereas no differences were detected at 3 or 6 months **(Fig. 5S, T; Suppl. Fig. 4A-C)**. Locomotor activity was unchanged, and FC confirmed intact associative memory across all timepoints **(Suppl. Fig. 4D-I)**. Overall, these data suggest that while LC tau pathology persists through 6-9 months, early noradrenergic hyperactivity and anxiogenic behaviors resolve over time, which is associated with reduced NE synthetic capacity^24^.

## Discussion

In MBI, NPS often emerge years before cognitive decline in AD, yet the mechanisms driving these behavioral manifestations remain elusive^3^. Here, we modeled Braak stage 0 by restricting pathogenic P364S human tau expression to the mouse LC, providing compelling evidence that LC-NE hyperactivity prior to LC degeneration drives mild behavioral phenotypes^9^. At 3 months post-infusion, Tau-expressing mice exhibited anxiety-and compulsive-like behaviors, diminished sleep spindle duration and density, and heightened spontaneous firing and excitability of LC neurons, with no change in LC cell bodies or fiber density. These functional alterations were accompanied by synaptic and metabolic pathway perturbations, positioning the LC as a critical node in early AD pathogenesis. Interestingly, anxiogenic phenotypes transitioned to anxiolysis at 6-9 months, suggesting a dynamic shift from noradrenergic hyperactivity to hypoactivity-associated states under long-term tau exposure^24^.

Although pathological tau typically suppresses neuronal activity, the LC-NE system engages compensatory mechanisms at multiple levels (cell bodies, dendrites, terminals, and postsynaptic receptors) to maintain homeostasis^29^. We speculate that these homeostatic plasticity processes “overshoot”, contributing to MBI in early AD. Consistent with this hypothesis, P364S tau markedly augmented both spontaneous and evoked LC firing at 3 months, aligning with observations from other models: (i) neuromelanin-accumulating noradrenergic neurons exhibit hyperactive properties^60^, (ii) pTau-positive LC neurons from young TgF344-AD rats show elevated activity in response to footshock^40^, and (iii) LC neurons in PS19 mice expressing brain-wide P301S human tau are hyperexcitable, with depolarized membrane potentials and increased spontaneous firing^41,69^ (but see^44^). This hyperactivity also mirrors early clinical AD, which is associated with increased TH expression and elevated CSF NE and metabolites^17,21–24^. Importantly, we observed no changes in NET expression or tau spread beyond the LC, indicating that compensatory changes in our model are primarily LC-intrinsic and cell-autonomous^26,28,32,34,56^. Furthermore, the striking astrocytic response within the LC parallels neuromelanin-based neurodegeneration models, suggesting that astrocytes are highly sensitive to LC-NE insults^60^.

Anxiety and agitation are recognized aspects of MBI, often manifesting as heightened vigilance and repetitive coping behaviors^70^. In affective disorders such as post-traumatic stress disorder, enhanced LC MRI signal correlates with symptom severity, implicating LC-NE hyperactivity as a key contributor of pathological anxiety^71,72^. Consistent with this framework, Tau-expressing mice exhibited greater stress-induced NS and prolonged feeding latencies in a novel environment at 3 months.

Beyond affective changes, LC firing varies as a function of behavioral states, including sleep stages^6,73^. Although overall sleep architecture remained intact, Tau-expressing animals showed a dramatic decrease in sleep spindles, consistent with LC hyperactivity; optogenetic stimulation of the LC destabilizes and suppresses spindle dynamics, promoting microarousals that interfere with memory consolidation and compromise restorative NREM functions, including glymphatic metabolic clearance^74–77^. These findings are consistent with prior work in PS19 mice, which display reduced spindle density at 2 months, along with significant alterations in REM duration and increased sleep fragmentation later in life^50^. However, the presence of ubiquitous pTau in PS19 mice may account for the absence of macrostructural sleep changes in our LC-restricted model^50^. Notably, in cognitively normal older adults, reduced spindle density and coupling to slow oscillations are associated with CSF total tau and tau-PET burden, supporting our interpretation that pathological tau disrupts sleep-related microstructural processes.

As Braak stages advance, LC volume declines, accompanied by frank neuronal degeneration^12,13^. In PS19 mice, LC lesions exacerbate hippocampal neurodegeneration and impair hippocampal-dependent associative memory, while 16 month-old TgF344-AD rats display progressive reductions in NE content, reduced stimulus-evoked LC firing and deficits in reversal spatial learning that are rescued by chemogenetic LC stimulation^33,34,40^. To determine whether prolonged accumulation of P364S human tau in the LC induces NE deficiency-associated hypoactive phenotypes, we aged animals 6 and 9 months post-infusions. Interestingly, the affect-related behavioral phenotypes observed at 3 months were absent. Moreover, Tau-expressing animals buried fewer marbles at 9 months, indicative of anxiolytic responses. TH IR was significantly reduced at 6 months and trended downward at 9 months, despite preserved LC cell number; similarly, no LC cell death has been reported in aged PS19 mice^33^. This pattern mirrors human pathology, in which LC neurons can harbor pathological tau for decades before degeneration; AT8 IR appears at Braak stage 0, but LC cell death does not occur until Braak stage III^11,13,27^. Furthermore, tau pathology assessed by AT8 and PHF1 IR remained unchanged across timepoints, suggested that longer pTau expression or fibrillar tau species may be required to elicit frank LC degeneration.

Clinical investigations of LC molecular alterations remain limited and largely rely on postmortem tissue, constraining insight into early disease dynamics. Given the pronounced phenotypes at 3 months, we focused on this timepoint. Changes associated with LC hyperactivity were evident, including upregulation of the hyperpolarization-gated channel *Hcn2*, which contributes to pacemaker currents and neuronal excitability, and is similarly increased by chronic morphine exposure in the ventral tegmental area, enhancing firing probability of dopamine neurons^78^. We also observed downregulation of the *Clic6* chloride channel, potentially disrupting inhibitory ion flux^79^. The upregulation of *Adra2c*, despite 𝛼_2A_ mediating ∼90% autoreceptor inhibition under physiological conditions, may suggest a compensatory response to pTau-induced hyperexcitability^80^. Additional DEGs included *Tesk1*, which regulates microtubule dynamics, and *Dapk3*, previously validated as a tau-interacting kinase in AD brain lysates^81,82^.

Tau-upregulated genes were enriched in WNT and CREB signaling and Class B2 GPCR pathways, suggesting compensatory mechanisms aimed at mitigating tau-induced cellular stress^83–85^. Downregulated pathways were dominated by synaptic and cytoskeletal programs. These findings parallel a recent single-nucleus RNA-sequencing study of noradrenergic neurons expressing pseudophosphorylated human tau, which reported convergent disruption of mitochondrial and synaptic transcripts despite minimal DEG overlap with our data^86^. Transcription factor analysis further indicated dampening of Atf4-driven transcription, a regulator of unfolded protein responses, and Ybx1, which controls DNA damage responses and RNA stress granules^87,88^. Moreover, Tau appears to disrupt cellular identity-associated programs, evidenced by Pax7 dysregulation (which governs pontine identity), Hes1 de-repression, and a shift towards Atf3 injury-driven signaling pathways^63,89,90^. Collectively, the translatome highlights a complex regulatory program that pTau-burdened LC neurons engage with respect to hyperexcitability, compensatory signaling, and stress adaptation.

We extended our analysis to the proteome. To overcome limitations in cellular specificity with bulk tissue proteomics, we adapted the CIBOP technique, enabling isolation of 862 high-confidence proteins from TH^+^ LC neurons. Critically, our dataset was enriched for canonical noradrenergic markers, including TH, DBH, NET, DDC and VMAT2, validating this approach. Proteomic profiling revealed a neuronal signature with enrichment in synaptic and neuronal compartments, cellular projections, and chemical transmission pathways. A recent study using DBH-Cre × Ai14 tdTomato mice isolated LC neurons to examine sex differences in the proteome and shared 52 proteins with our dataset (data not shown); however, their single-cell proteomics did not capture canonical LC-NE markers, underscoring the enhanced specificity and sensitivity of the CIBOP method^91^.

Analysis of pTau-induced proteomic changes revealed several notable and novel alterations. SNAP47, a synaptic machinery protein whose reduction is associated with cognitive decline in AD, was unexpectedly increased, indicating compensatory synaptic mechanisms during early disease stages^92^. UBR3, an E3 ubiquitin ligase involved in protein degradation, and RAB3IP, previously shown to rise in response to neurotoxic insults in PD models, were both significantly elevated^68,93^. CSNK1G2, a casein kinase phosphorylating diverse substrates is relatively unexplored in AD and was upregulated at both the mRNA and protein level^94^. We also observed enrichment of mTOR, which promotes tau hyperphosphorylation and accumulation^95^. Together, these changes map to pathways associated with signal transduction, catalytic activity, and increased metabolic demand, including enrichment of carbohydrate-derivative metabolic processes. Concomitant reductions in TCP1, CCT2, and CCT3, components of the chaperonin complex essential for cytosolic protein folding and tau aggregate clearance, were detected, consistent with tau-driven proteostatic and structural instability observed in our transcriptomic analysis^96^. Comparison with human cortical network modules indicates that LC tau engages similar functional systems (protein trafficking and neurotransmitter regulation), but display region-specific organization, consistent with LC-specific remodeling rather than cortical network replication^97^.

### Caveats and considerations

Some limitations should be considered. The P364S variant is an FTD-associated mutation that does not cause AD, though prior work indicates that it induces tau hyperphosphorylation and aggregation which resembles AD pathology^45,46^. While pTau accumulation was primarily restricted to the LC, trace ectopic expression was detected in the neighboring parabrachial nucleus, consistent with known Cre expression patterns in TH-Cre mice^98^. Spatial variability in viral expression within the LC could not be directly verified in omics-based samples.

We also noted limited concordance between Tau-induced mRNA and protein changes, which reflects known differences in RNA-protein stability, half-life, and/or subcellular localization^99^. The transcriptome captured ion channel dynamics, but the TurboID-based approach is biased toward cytosolic proteome with may underrepresent plasma membrane-bound proteins, likely due to the nuclear export sequence tag^49^. Finally, neuronal activity, sleep architecture, and molecular alterations were primarily assessed at the 3-month timepoint, limiting inference about later-stage molecular dynamics.

### Conclusions

Our findings provide evidence that pTau accumulation in the mouse LC is sufficient to induce neuronal hyperactivity and contributes to anxiety-like behavioral phenotypes during early disease stages. This work represents the first *in vivo* characterization of the P364S variant and integrates transcriptomic and proteomic analyses to define early molecular adaptations within this small brainstem nucleus. Furthermore, the application of CIBOP demonstrates the feasibility of LC-specific proteomic profiling, opening avenues to investigate mechanistic questions in neurodegeneration and beyond. Future studies can extend these findings by examining long-term pathogenic tau effects on LC-NE integrity and transmission.

## Acknowledgements

This research project was supported in part by the Viral Vector Core of the Emory Center for Neurodegenerative Disease Core Facilities. We thank Dr. Maureen Sampson for the guidance provided on transcriptomic data management. We also thank the MS & Proteomics Resource at Yale University for providing expertise on MS experiments, funded in part by the Yale School of Medicine and by the Office of the Director, National Institutes of Health (S10OD02365101A1, S10OD019967, and S10OD018034). The funders had no role in study design, data collection and analysis, decision to publish, or preparation of the manuscript. We thank Ms. Hailian Xiao and Mr. Lihong Cheng for support for animal breeding and husbandry.

## Author contributions

A.K. and D.W. conceptualized the study, developed the methodology, and wrote the original draft.

A.K., H.E.B., K.K., D.K., C.E., W.E.J., M.M.T., R.W., C.C.R., B.S.P., L.C.L., L.L., J.A., and E.G. conducted the investigation. Formal analysis was performed by A.K., H.E.B., K.K., U.S., and B.M. Data curation was carried out by A.K., U.S., and B.M. Validation and visualization were performed by A.K. Writing, review, and editing were carried out by A.K., H.E.B., K.K., U.S., B.M., A.W.V., S.R., M.J.B., and D.W. Supervision was provided by K.M., A.L.S., A.W.V., S.R., M.J.B., and D.W. Funding was acquired by A.K., H.E.B., A.L.S., A.W.V., S.R., and M.J.B.

## Data Availability

All sequencing data generated in this study are available in the NCBI Gene Expression Omnibus (GEO) under accession number **GSE314950**. Proteomics mass spectrometry data are available via the PRIDE repository under accession number **PXD071656**. Additional datasets generated during this study are available from the corresponding author upon reasonable request.

## Ethics Statement

All animal procedures were approved by the Institutional Animal Care and Use Committees (IACUCs) of Emory University, the Oklahoma Medical Research Foundation, and the Icahn School of Medicine at Mount Sinai and conducted in accordance with NIH guidelines.

