## Supplemental Material for "Pathogenic tau in the mouse locus coeruleus produces noradrenergic hyperactivity and neuropsychiatric phenotypes reminiscent of early Alzheimer’s disease"

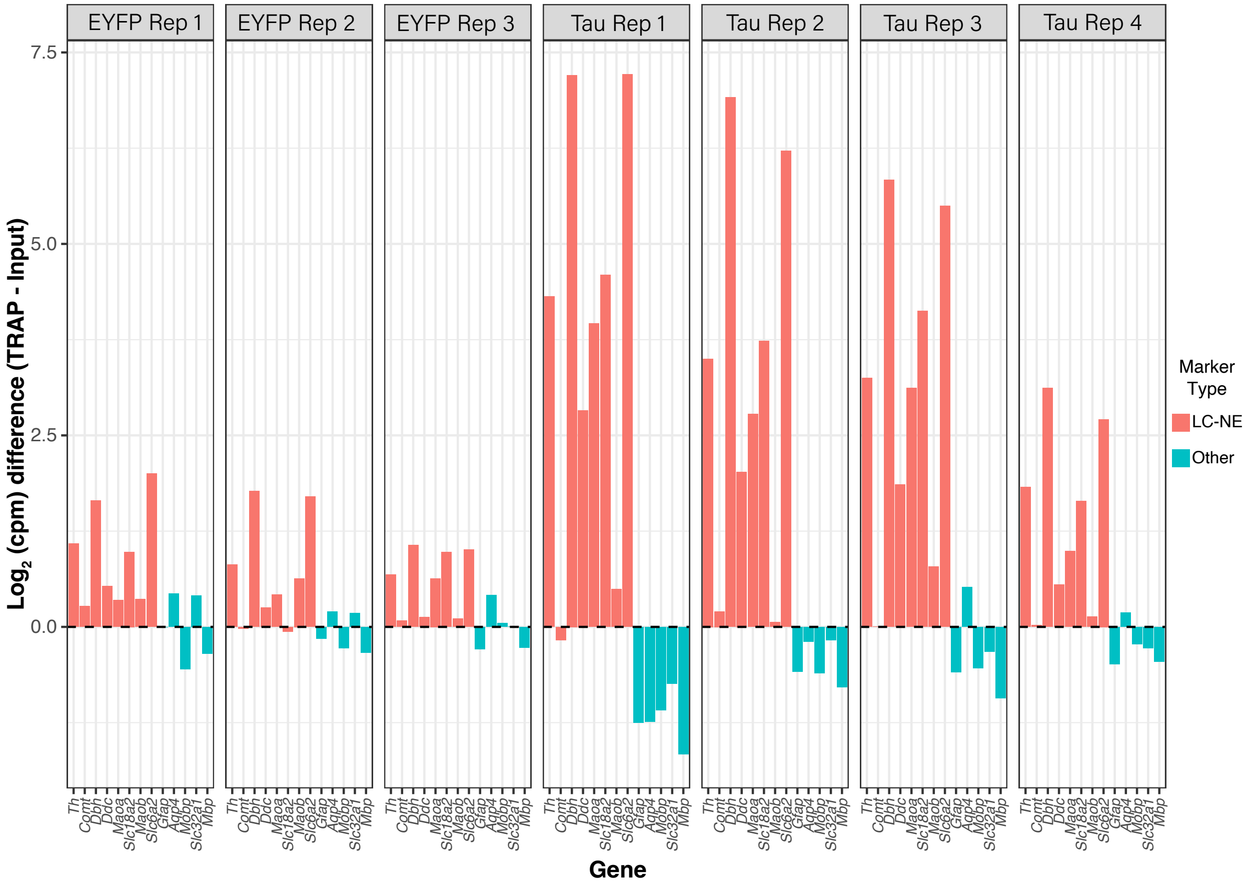

EYFP Rep 1
EYFP Rep 2
EYFP Rep 3
Tau Rep 1
Tau Rep 2
Tau Rep 3
Tau Rep 4
7.5
5.0
2.5
0.0
Log2 (cpm) difference (TRAP - Input)
Th
Comt
Dbh
Ddc
Maoa
Slc18a2
Maob
Slc6a2
Gfap
Aqp4
Mobp
Slc32a1
Mbp
Gene
Marker Type
LC-NE
Other


585

**Supplementary Figure 1. Enrichment of LC-NE genes and other cell-type markers in the IP fraction of TRAP samples.**

TRAP efficiency for each sample was evaluated by calculating the $log_2$ counts per million (cpm) difference between TRAP IP and Input fractions for known cell-type markers. Markers included LC-NE (*Th, Comt, Dbh, Ddc, Maoa, Slc18a2, Maob, Slc6a2*), astrocytes (*Gfap, Aqp4*), oligodendrocytes (*Mobp, Mbp*), and GABA (*Slc32a1*).

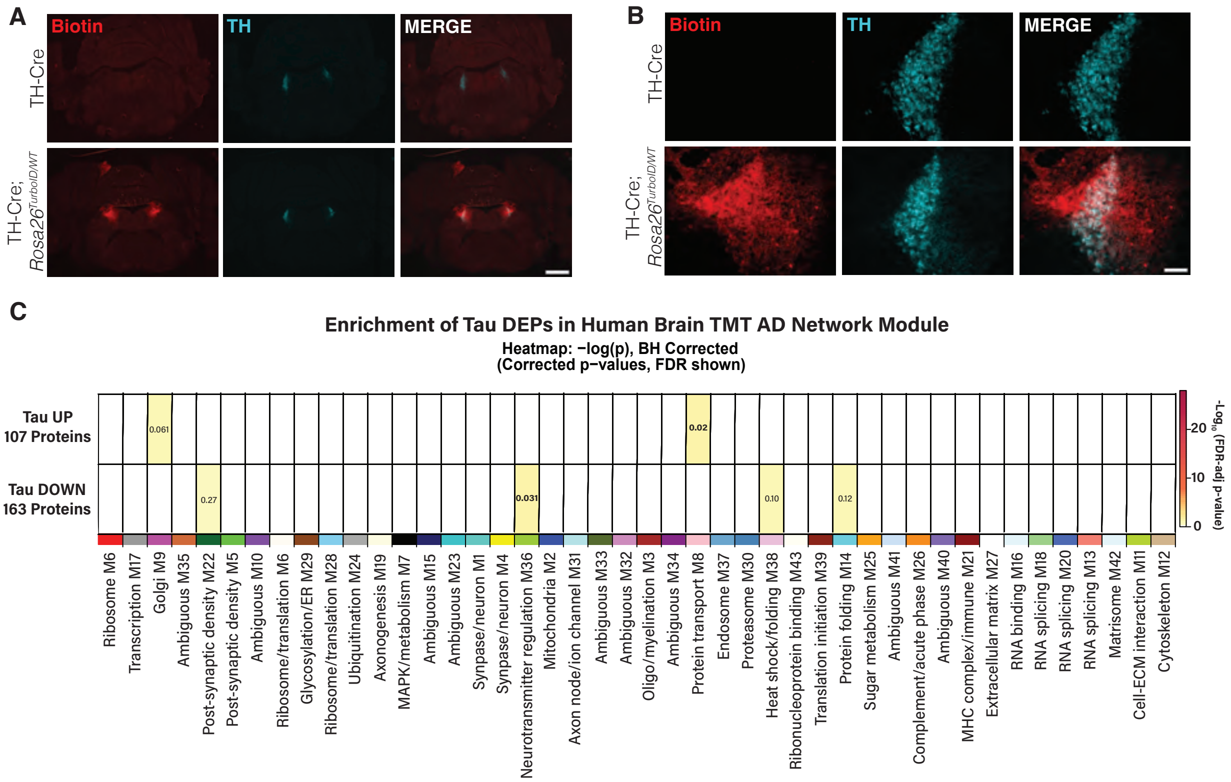
A
Biotin
TH
MERGE
TH-Cre
TH-Cre;
Rosa26TurboID/WT
B
Biotin
TH
MERGE
TH-Cre
TH-Cre;
Rosa26TurboID/WT
C
Enrichment of Tau DEPs in Human Brain TMT AD Network Module
Heatmap: −log(p), BH Corrected
(Corrected p−values, FDR shown)
Tau UP
107 Proteins
Tau DOWN
163 Proteins
0.061
0.02
0.27
0.031
0.10
0.12
-Log10 (FDR-adj p-value)
20
10
0
Ribosome M6
Transcription M17
Golgi M9
Ambiguous M35
Post-synaptic density M22
Post-synaptic density M5
Ambiguous M10
Ribosome/translation M6
Glycosylation/ER M29
Ribosome/translation M28
Ubiquitination M24
Axonogenesis M19
MAPK/metabolism M7
Ambiguous M15
Ambiguous M23
Synpase/neuron M1
Synpase/neuron M4
Neurotransmitter regulation M36
Mitochondria M2
Axon node/ion channel M31
Ambiguous M33
Ambiguous M32
Oligo/myelination M3
Ambiguous M34
Protein transport M8
Endosome M37
Proteasome M30
Heat shock/folding M38
Ribonucleoprotein binding M43
Translation initiation M39
Protein folding M14
Sugar metabolism M25
Ambiguous M41
Complement/acute phase M26
Ambiguous M40
MHC complex/immune M21
Extracellular matrix M27
RNA binding M16
RNA splicing M18
RNA splicing M20
RNA splicing M13
Matrisome M42
Cell-ECM interaction M11
Cytoskeleton M12

**Supplementary Figure 2. Validation of CIBOP model for cell type-specificity.**

Representative IF images showing colocalization of biotin (red) in $TH^+$ cells (cyan) at **(A)** 2x, scale bar = 1000μm and **(B)** 20x, scale bar = 100μm. **(C)** Heatmap showing overlap between tau-regulated DEP sets and 44 human brain protein co-expression modules from the AMP-AD Consortium proteomic dataset. Modules were defined using WGCNA from deep tandem mass tag mass spectrometry (TMT-MS) profiling of postmortem human brain tissue. Enrichment was assessed using Fisher's exact tests with FDR correction. Color intensity represents $-Log_{10}$ (FDR-adjusted p-value).

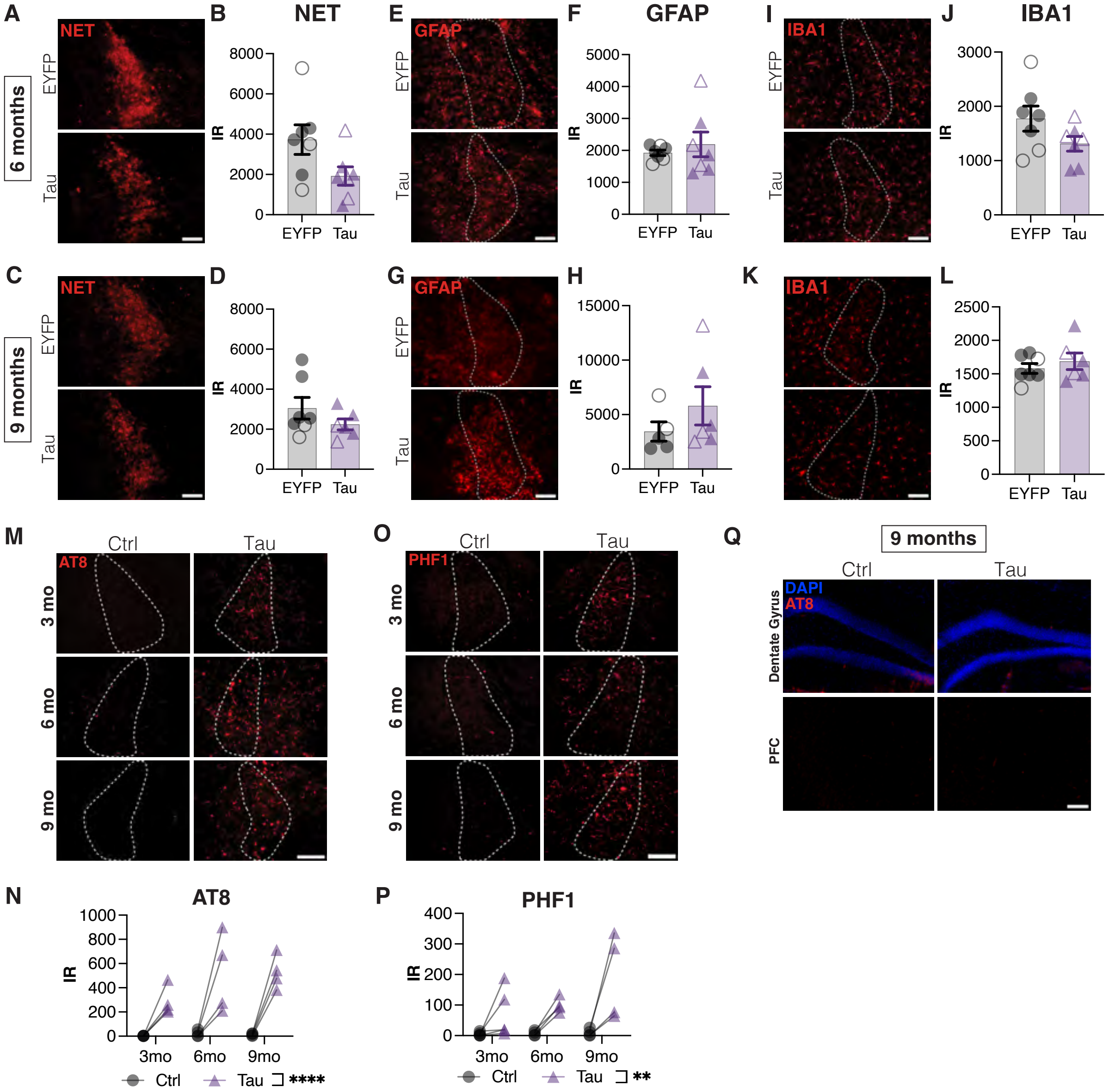
A
NET
6 months
EYFP
Tau
B
NET
IR
8000
6000
4000
2000
0
EYFP
Tau
E
GFAP
F
GFAP
5000
4000
3000
2000
1000
I
IBA1
J
IBA1
3000
2000
1000
C
9 months
D
G
H
15000
10000
5000
K
L
2500
2000
1500
1000
500
M
Ctrl
AT8
3 mo
6 mo
9 mo
O
PHF1
Q
9 months
DAPI
Dentate Gyrus
PFC
N
AT8
800
600
400
200
3mo
6mo
9mo
Ctrl
Tau ****
P
PHF1
400
300
200
100
Tau **

**Supplementary Figure 3. Prolonged tau burden does not affect LC integrity or neuroinflammation at 6 or 9 months.**

Representative images and quantification of IR of **(A-D)** NET (red), **(E-H)** GFAP (red), **(I-L)** IBA1 (red) (LC outlined in white; 20x, scale bar = 100 μm) at 6 and 9 months (n = 7 per group; unpaired t-test). Representative images and quantification of **(M, N)** AT8 and **(O, P)** PHF1 through 3, 6, and 9 months post-infusions (n = 4 per group, two-way repeated-measures ANOVA). **(Q)** Representative images of AT8 IR (red) in the DG and PFC at 9 months. Data are presented as mean ± SEM (male: closed, female: open symbols). **p = 0.001, ****p < 0.0001.

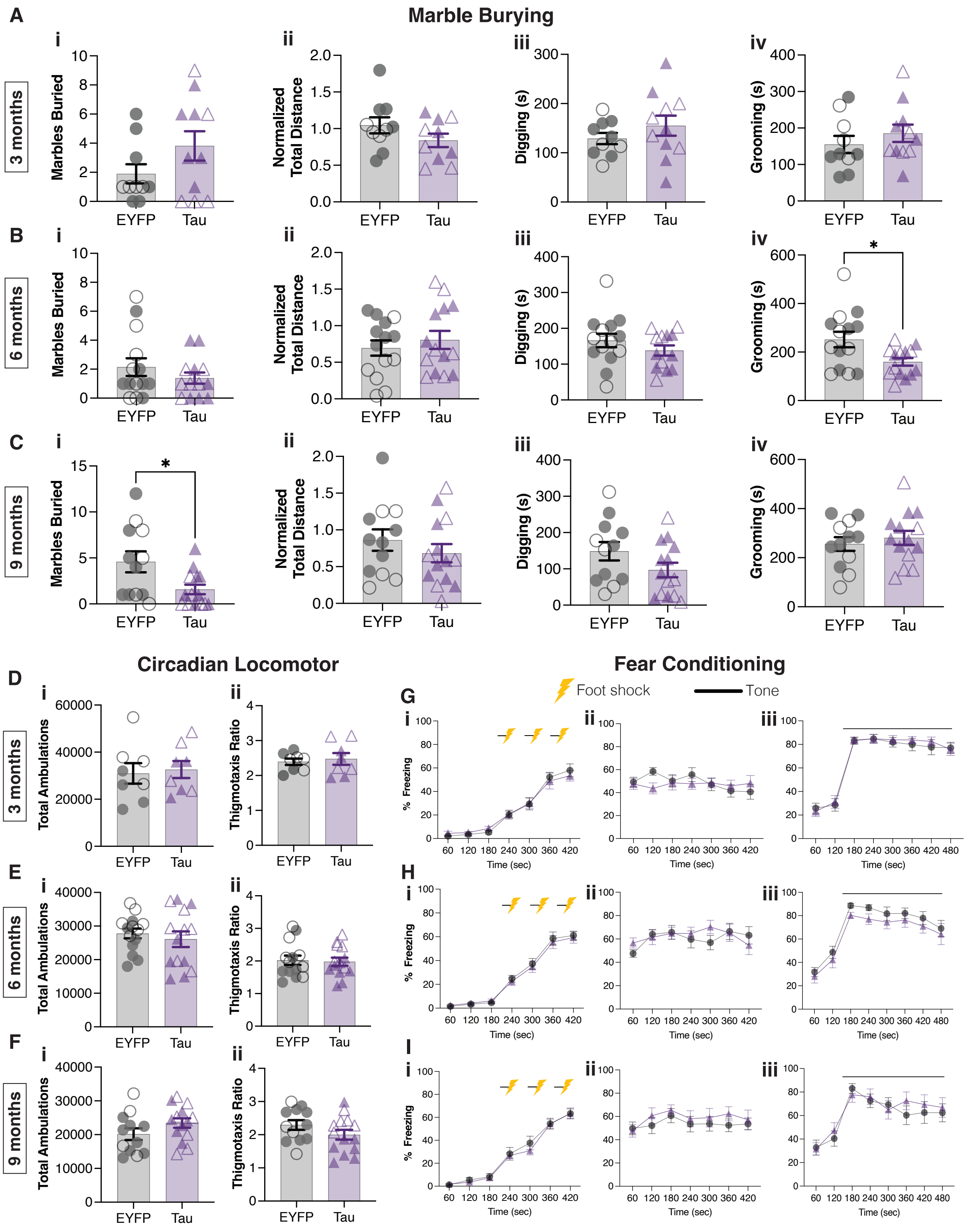

Marble Burying
A
3 months
6 months
9 months
Marbles Buried
Normalized Total Distance
Digging (s)
Grooming (s)
EYFP
Tau
B
C
Circadian Locomotor
D
E
F
Total Ambulations
Thigmotaxis Ratio
Fear Conditioning
Foot shock
Tone
G
H
I
% Freezing
Time (sec)

**Supplementary Figure 4. Pathogenic tau in the LC does not alter locomotor activity or associative memory.**

**(A-C)** Bar graph showing the (i) number of marbles buried, (ii) normalized total distance covered (iii) time spent digging (iv) time spent grooming. **(D-F)** Bar graph displaying (i) locomotor activity and (ii) thigmotaxis ratio (A-D: unpaired t-test) **(G-I)** Line graph showing fear behavior as a percentage of time spent freezing on (i) training day (ii) contextual memory (iii) cued memory (two-way ANOVA). Data are presented as mean ± SEM (n = 10-14 per group; male: closed, female: open symbols). *$p < 0.05$.

### Supplementary Methods

**Table 1. List of antibodies used for immunofluorescence and western blots.**

| Antibodies | Host | Manufacturer | Catalog No. | Dilution |
|---|---|---|---|---|
| **Immunofluorescence** | | | | |
| Tyrosine hydroxylase (TH) | Chicken | Abcam | Ab76442 | 1:1000 |
| Norepinephrine transporter (NET) | Mouse | Mab Technologies | NET05-2 | 1:1000 |
| Human Tau | Mouse | Synaptic Systems | 314 111 | 1:1000 |
| Phospho-Tau (Ser202, Thr205) AT8 | Mouse | Invitrogen | MN1020 | 1:1000 |
| Phospho-Tau (Ser 396/404) PHF1 | Mouse | Dr. Peter Davies (Albert Einstein College of Medicine) | - | 1:1000 |
| Glial fibrillary acidic protein (GFAP) | Rabbit | Abcam | Ab7260 | 1:1000 |
| Ionized calcium-binding adaptor molecule 1 (IBA1) | Rabbit | FUJIFILM Wako Pure Chemical Corporation | 019-19741 | 1:1000 |
| Alexa Fluor 568 Anti-Mouse | Goat | ThermoFisher Scientific | A11004 | 1:500 |
| Alexa Fluor 568 Anti-Rabbit | Goat | ThermoFisher Scientific | A11011 | 1:500 |
| Alexa Fluor 647 Anti-Chicken | Goat | ThermoFisher Scientific | A21449 | 1:500 |
| Streptavidin, DyLight 594 | | Invitrogen | 21842 | 1:500 |
| **Western Blot** | | | | |
| Tyrosine hydroxylase (TH) | Mouse | Immunostar | 22941 | 1:10,000 |
| Tau (phospho T231) antibody [EPR2488] | Rabbit | Abcam | ab151559 | 1:1000 |
| β-Actin | Mouse | Sigma Aldrich | A3854 | 1:20,000 |
| Streptavidin, Alexa Fluor 680 | | Invitrogen | S32358 | 1:10,000 |

**Behavioral paradigms**

<u>Nestlet shredding</u>: Mice were placed into a novel standard mouse cage (13” x 7” x 6”) with clean bedding and a pre-weighed cotton nestlet square (5 cm x 5 cm, ~ 3 g) and left undisturbed for 4 h. Nestlet weights were recorded at 2 h and 4 h[100].

Marble burying: Mice were placed into a large novel arena (24” x 12” x 12”) with 20 black glass marbles arranged in a 4 x 5 grid pattern on top of 2” of clean standard bedding and undisturbed for 30 min. The number of marbles buried, defined as ≥ 2/3 of marble being submerged in the bedding[100]. To quantify marble distance travelled, marble positions were tracked at fixed intervals during testing. At trial onset (t = 0), mice were placed into the cage, and images were acquired every 5 min to record marble positions. Total standardized distance traveled was calculated as the sum of individual movement distances normalized to the x-axis length. Grooming was defined as repetitive paw movements directed toward the whiskers and face, as well as licking or scratching of the body and tail. Digging presented as intense burrowing involving substrate displacement, resembling “swimming” or tunneling through bedding. Grooming and digging durations were quantified by video analysis, recording behavior to the nearest second.

Novelty-Suppressed Feeding: Following 24 h food deprivation, mice were habituated to the testing room for 2 h and then placed in a novel arena (24” x 12” x 12”) of a thin layer of standard bedding substrate with a single food pellet in the center. The latency for the mouse to feed (grasp and bite the food pellet) was recorded. Mice that did not feed within 15 min were assigned a latency score of 900 sec. Testing was conducted under red light to reduce the effect of bright light on anxiety-like responses. To eliminate hunger as a confounding variable, the test was repeated in the home cage.

Fear conditioning: Mice were placed in a fear-conditioning chamber (7 in x 7 in x 12 in H10-11M-TC, Coulbourn Instruments) equipped with a house light, ceiling-mounted camera, speaker, and metal shock grid floor, which could be replaced with nonshock wire mesh floor. On Day 1, the acquisition trial lasted for 7 min and consisted of 3 min acclimatization period followed by 3 tone-shock pairings with a 1 min inter-trial period. The conditioned stimulus was a 20 sec, 85 dB tone which co-terminated with the unconditioned stimulus 0.5 mA footshock (Precision Animal Shocker, Colbourn Instruments). On Day 2, contextual fear testing was conducted in the same chamber as Day 1 without any tone-shock presentation and continued for 8 min. On Day 3, cued fear testing was performed in a different chamber than Day 1 or 2 and the grid floor was replaced with the wire mesh. The trial lasted for 9 min, with tone presentation starting at 3 min and lasting till the end of the trial. Freezing behavior was recorded using the FreezeFrame software (Coulbourn Instruments) during each trial as a proxy for associative memory.

**Sleep studies**

Recordings were conducted in the home cage with a suspended multichannel commutator (Pinnacle Technologies). Signals were acquired at 1000 Hz sampling rate and bandpass filtered from 0.5–100 Hz (Pinnacle). The 24-h video recordings were collected at 10 frames/sec (synchronized with EEG) using an infrared LED camera with Sirenia Video Acquisition software (Pinnacle) and analyzed in Matlab with the Fieldtrip toolbox.

Behavioral states (rapid eye movement (REM), non-REM (NREM), wakefulness) were automatically scored based on previous criteria with manually curated review (see Supplementary Methods for more information). Sleep architecture was assessed by quantifying active and quiet wakefulness, REM, and NREM. Arousal, defined as wake bouts that occurred after REM or NREM and last for 1-60 seconds, was used to determine the arousal index (number of arousals divided by total sleep time). Sleep spindles and slow oscillations were detected as previously described[50,101].

**Electrophysiology**

For slice preparation, horizontal brain slices at the level of the fourth ventricle were placed in a holding chamber containing aCSF (in mM, 126 NaCl, 2.5 KCl, 1.2 $MgCl_2$, 2.4 $CaCl_2$, 1.2

$NaH_2PO_4$, 21.4 $NaHCO_3$, and 11.1 glucose) supplemented with 1 Na-ascorbate, 1 Na-pyruvate, 6 N-acetyl-L-cysteine, and 0.01 MK-801 for 30 min at 32 °C prior to recording. Neurons were recorded up to eight hours after slicing.

Following recovery, brain slices were transferred to a recording chamber and perfused with aCSF (32-34 °C, Warner Instruments inline heater) at a rate of 2 mL/min (Warner Instruments peristaltic pump). Individual LC neurons were identified by 1) location: rostral to the fourth ventricle and mediolateral to the large mesencephalic trigeminal tract (Me5) neurons and cytoarchitecture, large >20µm cell bodies, 2) electrophysiological properties: slow (0.5-5Hz) autonomous action potential generation in the cell attached configuration, steep input-output relationship to somatic current injection, and a slow (decay tau >75ms) A-type potassium conductance. Cell-attached and whole cell recordings were made with thin-wall glass (World Precision Instruments or Warner Instruments) pulled to a tip resistance of 2.5-3 MΩ and 2-2.5 MΩ, respectively. All recordings were made with an internal recording solution containing (in mM) 135 K-gluconate, 10 HEPES, 5 KCl, 5 $MgCl_2$, 0.1 EGTA, 0.075 $CaCl_2$, 2 ATP, 0.4 GTP, pH 7.35, which has a calculated liquid junction potential of -17.1mV that was not corrected (https://swharden.com/LJPcalc). Recordings were acquired via a Multiclamp 700B amplifier (Molecular Devices) and digitized with an Instrutech ITC-18 board. Data were acquired using AxoGraph (v1.8.0). All recordings were digitized at 20kHz. Current clamp recordings were low-pass filtered at 10kHz, and cell-attached voltage clamp recordings were low-pass filtered at 6kHz. All data were batch-analyzed off-line using custom written Python routines.

**TRAP and RNA sequencing**

RNA was extracted using Zymo RNA Clean & Concentrator-5 kit. Sample quality was assessed by High Sensitivity RNA Tapestation (Agilent Technologies Inc., California, USA) and quantified by AccuBlue Broad Range RNA Quantitation assay (Biotium, California, USA). Library construction was performed with SMART-Seq v4 Ultra Low Input RNA Kit (Takara Bio USA Inc., California, USA) followed by Nextera XT DNA Library Prep Kit (Illumina, California, USA). Final library quantity was measured by KAPA SYBR FAST qPCR and library quality evaluated by TapeStation D1000 ScreenTape (Agilent Technologies, CA, USA). Final library size was about 430bp with an insert size of about 200bp. Illumina 8-nt dual-indices were used. Equimolar pooling of libraries was performed based on QC values and sequenced on an Illumina® NovaSeq X plus (Illumina, California, USA) with a read length configuration of 150 PE for 40 M PE reads per sample (20M in each direction). Fastq files were assessed for quality using FastQC and Nextera adaptor trimming was performed using trimGalore. Reads were mapped to the mouse genome GRCm38 (mm10) and human tau using STAR alignment and raw counts were obtained using the featureCounts package in R Bioconductor.

**Differential gene expression analysis for TRAP**

Gene counts data were imported into R 4.4.3 on a Macintosh for analysis using *edgeR*'s *voomLmFit*. Counts from replicates where both a TRAP and Input (total homogenate RNA) were collected. We first checked for successful enrichment of LC marker genes by TRAP, which revealed a systematic group difference in enrichment efficiency. We therefore only analyzed replicates from which both a TRAP IP and Input fraction had been sequenced. To perform differentially expressed gene (DEG) analysis, counts data only from those replicates were imported and distributions of counts visualized to determine a low-count cutoff, which we set to 89, requiring at least 4 TRAP samples to exceed this threshold for a given gene. 14,224 genes and the hTauP364S transgene were therefore retained for downstream differential expression analyses. TMM normalization factors were then calculated to account for sequencing library size[102].

TRAP efficiency normalization factors were calculated for each pair of TRAP IP and Input by computing the $\log_2$ counts per million (cpm) of *Slc6a2*, *Dbh*, and *Th*. The mean $\log_2$(CPM) of these three genes were then obtained for each IP and Input, and the difference in these values between TRAP and Input was used as the enrichment adjustment factor for downstream DEG analyses. As sexes were also imbalanced between the two conditions after subsetting to samples with both a TRAP and Input fraction sequenced, we additionally controlled for sex in the DEG model. The final model assessed only TRAP fractions, under a model of *~0+transgene+sex+TRAPefficiency*. *voomLmFit* was run without sample weights to avoid interfering with the sample-specific, continuous adjustment variable already present (*i.e.*, the TRAP efficiency factor). We subsequently used *edgeR's topTable* to collect full DEG results.

**Gene Set Enrichment Analysis (GSEA) for TRAP**

To test for enrichment of DEGs in ontology and literature-derived sets, we utilized the threshold-free GSEA method [103] and its accompanying gene set catalogs (curated/C2, regulatory targets/C3, ontologies/C5, oncogenic/C6, and cell type signatures/C8) in mSigDB version 2024.1 mouse, implemented using the R package *fGSEA*[104]. We previously described our collation and use of several additional TF-target gene sets[60] from the *Enrichr* tool[105-107]: these include ChEA 2022[108], TRRUST[109], Rummagene publication mining[110], Enrichr-mined Gene Expression Omnibus TF perturbation experiments, and Enrichr's internal collection of TF-TF interactions[107]. We input the signed DE *t* statistics for use as the DEG ranking values to *fGSEA* with analysis parameters as follows: gene set sizes 15-500; *eps* (specifying range over which to calculate p-values) set to 0 to return a p-value for all tests; and 50,000 permutations for initial p-value estimations.

**Immunoblotting and affinity purification of biotinylated proteins**

To confirm protein biotinylation by western blot (WB), 10 μg of proteins from brainstem homogenates were separated on a 4-12% Bis-Tris gel (Invitrogen, NW04125BOX) at 80V and transferred to nitrocellulose membranes using the iBLOT mini stack system (Invitrogen, IB23002). After washing with 0.1% TBS-Tween 20, membrane was blocked with blocking buffer (ThermoFisher Scientific, 37543) for 1 h at room temperature (RT), probed with Streptavidin680 (Invitrogen, S32358) for 1 h at RT and then imaged on the ChemiDoc Imaging System (Bio-Rad). Next, membrane was reblotted with anti-pTau231 (Abcam, ab151559) and anti-TH (Immunostar, 22941) antibodies followed by corresponding secondary antibodies to validate treatment and cell-type specificity respectively. To confirm equal loading of the samples, immunoblots were probed with β-actin (Sigma-Aldrich, A3854).

Following previously optimized protocols[49,65], biotinylated proteins were captured by streptavidin magnetic beads (ThermoFisher Scientific, 88817), incubating 83 μL beads per 1 mg of protein in a 500 μL RIPA lysis buffer (50 mM Tris, 150 mM NaCl, 0.1% SDS, 0.5% sodium deoxycholate, 1% Triton X-100) for 1 h at 4˚C with rotation. After incubation, beads were then washed sequentially at RT 2x with RIPA buffer for 8 min/each, 1x with 1 M KCl for 8 min, 1x with 0.1 M $Na_2CO_3$, 1x with 2 M urea in 10mM Tris-HCl (pH 8.0) for 10 sec, 2x with RIPA lysis buffer for 8 min/each and 4x PBS. Finally, beads were then recovered on a magnetic rack, PBS was removed completely and further diluted in 100 μL PBS. For quality control studies by WB and silver staining, 10% of beads were eluted by heating the beads in 30 μL of 2X protein loading buffer (Bio-Rad, 1610737) supplemented with 2 mM biotin + 20 mM dithiothreitol at 95˚C for 10 min. Subsequently, 10 μL of eluate was run on a gel and probed with Streptavidin680 while 20 μL of eluate was run on a separate gel and Silver stained (ThermoFisher Scientific, 24612). The remaining 90% enriched biotinylated proteins were stored at -20˚C until on-bead digestion.

**On-Bead Digestion of Streptavidin-Enriched Pulldown Samples and Input Lysates**

On-bead digestion of proteins from streptavidin-enriched pulldown samples and digestion of input homogenates were performed using the S-Trap micro protocol by PROTIFI. Briefly, streptavidin bead-bound proteins and input protein homogenates (50 μg) were lysed in 5% SDS lysis buffer and sonicated for 10 minutes to ensure complete disruption. Samples were then incubated with 10 mM dithiothreitol (Sigma-ALDRICH #D5545) for 1 h at RT to reduce disulfide bonds. Following reduction, 30 mM iodoacetamide (IAA; Sigma-ALDRICH #I6125) was added for 30 minutes at RT in the dark to alkylate cysteine residues. To acidify the samples, 2.5% phosphoric acid was added, followed by addition of 76 mM triethylammonium bicarbonate (TEAB) pH 7.55 (Thermo Scientific #90114) binding buffer. Next, samples were loaded onto S-Trap micro columns (PROTIFI #C02-micro-80) placed in 2 mL collection tubes and centrifuged at 4,000×g for 30 seconds to bind proteins to the S-Trap matrix. After loading, the columns were washed 3 times with 100 mM TEAB (pH 7.55) buffer by centrifuging at 4,000×g for 1 minute each time. Proteins trapped in the columns were digested overnight with 1 μg of lysyl endopeptidase (Lys-C; Wako, #127-06621) at 37˚C. Next day, proteins were further digested using 2 μg trypsin (ThermoFisher Scientific, #90058) by overnight incubation at 37˚C. On the third day, peptides were eluted from the column using three different buffers sequentially: (1) 50 mM TEAB, (2) 0.2% formic acid, and (3) 50% acetonitrile. For each elution, addition of buffer was followed by centrifugation at 4000×g for 1 minute, and eluates were collected in fresh 1.5 mL tubes. The peptides were then dried using a vacuum concentrator (SpeedVac Vacuum Concentrator). Dried peptides were resuspended in loading buffer (0.1% formic acid), and peptide amount was measured by performing peptide BCA assay following manufacturer's protocol (Thermo fisher Pierce fluorometric kit #23290). Briefly, 10 μL of each standard (serially diluted, ranging from 500 pg to 7.8 pg) and sample was added into different wells of 96-well plate, followed by addition of 70 µL peptide assay buffer and 20 µL peptide assay reagent onto the samples and standards. Fluorescence was measured at 390 nm excitation and 475 nm emission.

**Mass spectrometry (MS)**

A MS analysis was performed in data-independent acquisition (DIA) mode using an m/z Range window mode. Samples for DIA were analyzed on a liquid chromatography (LC)–MS/MS consisting of a Vanquish Neo UHPLC (ThermoFisher Scientific) nanoflow LC system and an Orbitrap Exploris 480 mass spectrometer (ThermoFisher Scientific) equipped with an EASY-Spray ion sources. The MS dataset is available on PRIDE (PXD071656). For more details on the MS protocol, data processing and quantification, normalization parameters, differentially expressed proteins (DEPs), and Gene Ontology (GO) enrichment analysis, refer to Supplementary Methods.

**MS Protein Data Processing and Quantification**

Three hundred ng (inputs) or 10% (pulldowns) were loaded onto a trap column (300 µm inner diameters, 5 mm length, 5 µm C18 particles, PepMap Neo Trap, Thermo Fisher Scientific) before being separated on an EASY-spray HPLC analytical column (75 µm inner diameter, 60 cm length, 1.7 µm C18 particles, Aurora Frontier, Ionopticks) at a flow rate of 300 nL/minute. Mobile phase A and B were composed of HPLC water with 0.1% formic acid (FA), and 80% HPLC acetonitrile with 0.1% FA, respectively. The peptides were resolved by changing the gradient of mobile phase B as follows: 1% to 4% in 10 min, 4% to 25% in 60 min, and 25% to 65% in 45 min. The gradient was followed by a 20-minute wash step (65% to 99% in 1 min, hold at 99% for 19 min) and column re-equilibration. The voltage for electrospray ionization (ESI) was set to 2,100 V, and the ion transfer tube temperature was set to 275 °C. The DIA method consisted of one full MS1 scan followed by 60 DIA (MS2) scans. The MS1 scan range was set to 400–1000 m/z. MS1 scans were acquired at a resolution of 120,000 (at 200 m/z) with an RF Lens of 50%, a normalized AGC target of 300% (absolute target 3,000,000), and an 'Auto' maximum injection time. The 60 subsequent DIA (MS2) scans were acquired at a resolution of 30,000, with a normalized AGC target of 1000% (absolute target 1,000,000) and an 'Auto' maximum injection time. Fragmentation was induced by high-energy collision dissociation (HCD) at 28% of the normalized collision energy. The isolation window for precursor ions was set to 10 m/z with 1 m/z overlap, covering the precursor range m/z 400–1000. Internal calibration was carried out using the lock masses of plydimethylcyclosiloxane ions (m/z 445.12003) produced from ambient air.

The acquired DIA raw files were processed using DIA-NN (v2.2.0, Academia) in library-free mode. The search was conducted against a Mus musculus (Mouse) FASTA database (UniProt UP000000589, downloaded 2025.04.11) supplemented with common contaminants. Search parameters were set to Trypsin/P specificity, allowing for up to two missed cleavages. Carbamidomethylation (C) was set as a fixed modification. Variable modifications included N-terminal M excision, Oxidation (M), and N-terminal acetylation, with a maximum of three variable modifications per peptide. The peptide length was restricted to 7–30 amino acids, and the precursor charge range was set from 1 to 4. The analysis matched precursors within the m/z 400–1000 range and fragments within the m/z 150–2000 range. Mass tolerances for both MS1 and MS2 were set to 10 ppm, with a scan window of 15. The analysis used neural network (NN)-based, cross-validated machine learning and proteoform-level scoring. All results were filtered at a 1% precursor and protein group FDR. Protein quantification was performed using the QuantUMS strategy. Cross-run normalization was disabled within DIA-NN.

**Protein filtering, imputation, and normalization**

The input (whole proteome) and biotin-enriched IP fractions underwent different data preparation. In terms of the biotin-labeled experimental groups, proteins that were missing from ≥ 50% of the samples in each group were eliminated. Following filtering, 4579 proteins remained in the IP sample, and 6666 proteins were kept in the input sample out of the initial 7055 proteins identified. Subsequently, the samples were $log_2$-transformed and imputed in Perseus using default parameters (width = 0.3, downshift = 1.8) to replace low-abundance missing values. After that we applied column-sum median normalization to correct total intensity (protein loading) differences across samples and ensure uniformity between replicates.

**Differential protein expression analysis**

Differentially expressed proteins (DEPs) were identified by statistical testing of normalized and imputed samples. All relevant comparisons, which include TH-Cre;*Rosa26*$^{TurboID/WT}$ vs. WT (Input), TH-Cre;*Rosa26*$^{TurboID/WT}$ vs. WT (IP), P364S hTau vs. EYFP: TH-Cre;*Rosa26*$^{TurboID/WT}$ (Input), and P364S hTau vs Control (IP) were all evaluated by pairwise contrasts after a one-way ANOVA across sample groups. The following threshold was used ($p < 0.05$) to determine statistical significance. Proteins with an absolute $log_2$ fold change > 0 and $p < 0.05$ were considered DEPs and visualized in R using the ggplot2 packages.

**Functional and GO Enrichment Analysis**

Functional annotation and enrichment analysis of the significantly DEPs were performed using the cluster Profiler (v4.0) and Enrichr (v3.2) packages. Over-representation analysis was conducted across Gene Ontology (GO) categories, including Biological Process (BP), Molecular Function (MF), and Cellular Component (CC), using all proteins retained after filtering within each dataset (Input or IP) as background. Enrichment significance was determined based on an adjusted p-value < 0.05 following multiple testing correction. Visualization of enrichment profiles was carried out using ggplot2, where enrichment scores were represented as $-\log_{10}$ p values, and gene ratios were plotted to reflect the proportion of DEPs contributing to each GO term.

**Network Module Enrichment Analysis**

We used a published human brain proteomic reference dataset, generated through the AMP-AD Consortium, consisting of postmortem dorsolateral PFC brain tissue derived from the Religious Orders Study and Memory and Aging Project (ROSMAP, n = 84 control, 148 Asymptomatic AD and 108 AD) and the Banner Sun Health Research Institute (Banner, n = 26 control, 58 Asymtomatic AD and 92 AD), analyzed by deep tandem mass tag (TMT)-MS-based quantitative proteomics [97]. The dataset defines 44 protein co-expression modules identified by weighted gene co-expression network analysis (WGCNA), each functionally annotated and linked to AD-relevant biological processes. Using this framework, Tau-regulated DEP sets and modules were overlapped using Fisher's exact tests. FDR correction was applied to account for multiple testing. Enrichment results were visualized as a heatmap, with each cell representing the degree of overlap between a given human brain module and a tau-regulated protein set, with color intensity corresponding to -$\text{Log}_{10}$ (FDR-adjusted p-value).
